# UPF1 ATPase autoinhibition and activation modulate RNA binding kinetics and NMD efficiency

**DOI:** 10.1101/2023.11.03.565554

**Authors:** Joseph H. Chapman, Alice M. Youle, Acadia L. Grimme, Keir C. Neuman, J. Robert Hogg

## Abstract

The RNA helicase UPF1 interacts with mRNAs, mRNA decay machinery, and the terminating ribosome to promote nonsense-mediated mRNA decay (NMD). Structural and biochemical data have revealed that UPF1 exists in an enzymatically autoinhibited “closed” state. Upon binding the NMD protein UPF2, UPF1 undergoes an extensive conformational change into a more enzymatically active “open” state, which exhibits enhanced ATPase and helicase activity. However, mechanically deficient UPF1 mutants can support efficient NMD, bringing into question the roles of UPF1 enzymatic autoinhibition and activation in NMD. Here, we identify two additional important features of the activated open state: slower nucleic acid binding kinetics and enhanced ATP-stimulated nucleic acid dissociation kinetics. Computational modeling based on empirical measurements of UPF1, UPF2, and RNA interaction kinetics predicts that the majority of UPF1-RNA binding and dissociation events in cells occur independently of UPF2 binding. We find that UPF1 mutants with either reduced or accelerated dissociation from RNA have NMD defects, whereas UPF1 mutants that are more dependent on UPF2 for catalytic activity remain active on well-established NMD targets. These findings support a model in which the kinetics of UPF1-mRNA interactions are important determinants of cellular NMD efficiency.

## Introduction

Eukaryotic cells use mRNA decay to quickly regulate gene expression in response to changing conditions (1). In addition to allowing tight kinetic control of gene expression, mRNA decay provides a mechanism by which cells can mitigate the negative effects of genetic mutations or errors in mRNA biogenesis. A prototypical example of an mRNA decay pathway that uses a common set of factors to accomplish both transcriptome surveillance and gene expression regulation is nonsense-mediated mRNA decay (NMD) (2). Together, the targets of NMD have been estimated to compose 5-20% of human genes, and defects in NMD are associated with numerous genetic diseases and cancers (3, 4).

At the core of the NMD pathway is the RNA helicase UPF1, which non-specifically binds RNA (5–7) to act as a protein scaffold for other NMD and translation factors. These include UPF2 (8–10), the terminating ribosome (11–13), the RNA endonuclease SMG6 (14–18), and the RNase-recruiting heterodimer SMG5/SMG7 (14, 15, 18). As a superfamily 1 (SF1) helicase, UPF1 harnesses the chemical energy from ATP hydrolysis to translocate 5′-3′ on single-stranded RNA, unwind RNA secondary structures, displace proteins from RNA, and dissociate from RNA (19–23).

In the absence of UPF2, the autoinhibitory UPF1 N-terminal cysteine/histidine-rich (CH) domain docks onto the RecA2 domain, reducing ATPase activity. UPF2 relieves UPF1 autoinhibition by interrupting a conserved phenylalanine-isoleucine interaction (F192 and I757 in human UPF1; F131 and I693 in yeast UPF1) and rotating the CH domain 120° away from the RecA2 domain (10, 24). In this way, UPF2 binding converts UPF1 from the enzymatically autoinhibited “closed” state to an activated “open” state (10, 24). Correspondingly, mutation of UPF1 F192 to glutamate (F192E) biochemically substitutes for UPF2 binding, resulting in constitutively activated UPF1 (10, 24, 25). Despite extensive biochemical and structural studies, the mechanisms by which cellular NMD is affected by either UPF1 autoinhibition or activation by UPF2 remain largely unexplored.

In addition to enhancing UPF1 ATPase activity, UPF2 promotes UPF1 phosphorylation by the PI3K-like kinase SMG1 (12, 13, 26). Phosphorylated UPF1 in turn stimulates NMD by recruiting SMG6 and SMG5/SMG7 (15). Structural studies have identified a megadalton-scale SMG1 complex (SMG1C) consisting of SMG1, SMG8, and SMG9 bound to UPF1 and UPF2 (26–28). In this context, the UPF2-induced conformational swing of the CH domain has been proposed to facilitate positioning of UPF1 in the SMG1 kinase core (26) and subsequent release from SMG1C (27).

UPF2-induced undocking of the CH domain has also been proposed to reduce UPF1-RNA interactions. Recently, it was reported that UPF1-UPF2-RNA ternary complexes are unstable, due to competition between UPF2 and RNA for UPF1 (29). Moreover, the tendency of UPF1 to release RNA upon ATP hydrolysis implies that UPF2 may further disrupt UPF1-RNA interactions via stimulation of UPF1 ATPase activity (30). Consistent with these findings, the UPF1 F192E mutant reduced ATPase-dependent RNA binding in both *in vitro* assays (24) and cellular immunoprecipitation experiments (5). Enhancement of UPF1 ATPase activity by UPF2 has historically been viewed as a means of promoting decay, but these data imply that this mechanism may also be used to preserve RNA stability by enhancing UPF1 dissociation from potential NMD substrates.

The role of UPF1 autoinhibition and activation in the regulation of UPF1-RNA interactions is of particular interest in light of accumulating evidence that ATPase-stimulated dissociation is a critical mechanism to ensure efficient and accurate NMD. We recently identified UPF1 mutants that are poorly processive, unwind slowly, displace proteins from nucleic acid less efficiently, and have ATPase activity decoupled from unwinding but can still promote efficient NMD of canonical targets (31). These data, together with findings of differential rates of UPF1 release from NMD-sensitive and NMD-insensitive mRNAs (5, 32), point towards the ‘butterfly model’ of NMD, which proposes that UPF1 must efficiently dissociate from non-target RNAs but more stably associate with target RNAs to promote NMD. This model predicts that either enhanced or reduced rates of UPF1 ATPase-stimulated dissociation would negatively impact cellular NMD. Accelerated dissociation would reduce opportunities for UPF1 to productively assemble decay complexes in response to translation termination events (32–34). Conversely, slower UPF1 dissociation would prevent efficient sampling of the transcriptome, exemplified by ATPase-dead UPF1 mutants that lose decay target discrimination (5, 35).

Here, we use *in vitro* biochemical and biophysical assays, computational modeling, and cellular studies to understand the functional significance of the conformational change between the autoinhibited closed state and the activated open state of UPF1. We find that UPF1-UPF2 complexes and perpetually open UPF1 mutants have slower nucleic acid association kinetics and faster ATP-dependent nucleic acid dissociation kinetics compared to the closed state of UPF1. Computational kinetic modeling predicts that the effects of UPF2 on UPF1-RNA interactions are constrained by UPF2 abundance in cells, such that the majority of UPF1-RNA association and dissociation events occur in the absence of UPF2. Furthermore, we characterize previously known and new UPF1 mutants with varying degrees of autoinhibition to understand the biochemical and cellular consequences of switching between the closed and open states. We identify no cellular NMD defects in UPF1 mutants with increased reliance on UPF2 for activation but find that mutants that undergo either accelerated or slowed ATPase-stimulated dissociation have reduced NMD function.

## Materials & Methods

### Oligonucleotides and chemicals

All oligonucleotides were purchased from IDT, resuspended in 1XTE, and stored at −20°C. For pH 6.0 assays, ATP (Millipore Sigma) was dissolved in 5 mM MES pH 6.0 to 10 mM and stored at −80°C, then diluted to 5 mM in 5 mM MES pH 6.0 for day-of use through the plate reader injectors. For pH 7.5 assays, ATP and AMP-PNP were dissolved in 10 mM Tris-HCl pH 7.5 to 100 mM and stored at −80°C. Alexa Fluor maleimides were dissolved in DMSO to 10 mM and stored at −20°C protected from light.

### UPF1 and UPF2 protein expression and purification

UPF1-CH-HD and UPF1-CH expression and purification was performed as previously described with UPF1-HD (31). These constructs contained N-terminal 6xHis and calmodulin binding peptide tags. UPF2 761-1227 with a C-terminal CLIP tag synthesized by Twist BioScience did not express well, so the UPF2 761-1227 fragment (UPF2-S) was PCR amplified out using Phusion (primers listed in **Supplementary Table S1**) and ligated into the pET28 vector with an N-terminal 6xHis and calmodulin binding peptide tag. Expression and purification conditions were identical to that of UPF1-CH-HD, except the dialysis buffer contained 1.5X PBS pH 7.4 (KD Medical), 10% glycerol, and 2 mM DTT. UPF1-CH-HD dialysis buffer contained 1.5X PBS pH 7.4 (KD Medical), 20% glycerol, 1 µM ZnSO_4_, 0.66 mM MgOAc, 150 mM NaCl, 0.1% NP-40, and 1 mM DTT.

### In vitro transcription and substrate annealing for biochemical assays

*In vitro* transcription and annealing were performed as previously described (31). Oligonucleotide sequences are provided in **Supplementary Table S1**.

### UPF1 unwinding, dissociation, and equilibrium binding

Fluorescence-based unwinding assays (at 37°C), fluorescence anisotropy dissociation (FAD) assays (at 37°C), and equilibrium fluorescence anisotropy nucleic acid binding assays (at room temperature) were performed as previously described unless otherwise indicated (31).

Measurements with UPF2 were performed by incubating UPF2-S in 2-fold excess to UPF1-CH-HD for 10 min after the UPF1-CH-HD incubation with the substrate, unless otherwise stated. All assays were performed using a CLARIOstar Plus plate reader (BMG Labtech). Oligonucleotide sequences used for substrates are provided in **Supplementary Table S1**.

### Biolayer interferometry (BLI) to obtain UPF1 and UPF2 on- and off-rates

All buffers consisted of BLI buffer (20 mM Tris-HCl pH 7.5, 75 mM KOAc, 3 mM MgCL_2_, 1 mM DTT, and 0.02% Tween-20). Streptavidin-coated sensors (Sartorius Octet SA Biosensors) were pre-soaked in BLI buffer for at least 10 min before runs and dipped into wells of a microplate (greiner bio-one 655209) containing 200 µL BLI buffer for the indicated times. To tether substrates, sensors were dipped into wells containing BLI buffer supplemented with 10 nM of a dual-biotinylated DNA oligonucleotide (see **Supplementary Table S1**). To measure UPF1 on-rate, sensors were dipped into wells supplemented with the indicated concentrations of UPF1. To measure ATP-simulated dissociation of UPF1 from the substrate, sensors were dipped into wells with 2 mM ATP and 50 µM of trap strand. To measure UPF2 on-rate to UPF1, sensors containing DNA and UPF1 were dipped into wells supplemented with the indicated concentrations of UPF2. To measure UPF2 dissociation from UPF1, sensors were dipped into wells supplemented with 50 µM of trap strand.

### UPF1 and UPF2 fluorescent labeling using cysteine-maleimide chemistry

UPF1 and UPF2 proteins were dialyzed in DTT-free dialysis buffers for 4 hr at 4°C replacing with fresh buffer every hour. The UPF1 DTT-free dialysis buffer contained 1.5X PBS pH 7.4 (KD Medical), 1 µM ZnSO_4_, 0.66 mM MgOAc, and 150 mM NaCl. The UPF2 DTT-free dialysis buffer contained 1.5X PBS pH 7.4 (KD Medical) and 150 mM NaCl. After dialysis, 1 mM TCEP was added for 10 min on ice to reduce solvent-exposed cysteines. AlexaFluor maleimide stock was added to the indicated concentration and incubated at 4°C overnight to label proteins. To remove free dye, proteins were passed through 0.5 mL 40K MWCO Zeba Spin Columns three times that were washed with DTT-free dialysis buffer, then passed once through columns that were washed with dialysis buffer (see recipe above). Proteins were then aliquoted and flash frozen in liquid nitrogen.

### Microscale Thermophoresis (MST) for protein-protein binding measurements

All dilutions performed during microscale thermophoresis used Ligand Buffer containing 20 mM Tris-HCl pH 7.5, 75 mM KOAc, 3 mM MgCl_2_, 1 mM DTT, and 0.05% Tween-20. UPF2-S labeled with 10-fold molar excess of Alexa-Fluor 647 was diluted to a final concentration of 3.125 nM. UPF1 constructs were diluted to 4 µM in Ligand Buffer, and a two-fold serial dilution was performed to obtain sixteen concentrations of UPF1. Equal parts of UPF2 and UPF1 dilutions were mixed and loaded into Monolith Standard Treated Capillaries. MST was run using 50% LED and medium power, and curve fits were generated using the MO.Affinity Analysis Software from NanoTemper Technologies.

### In silico Rosetta-Vienna RNP ΔΔG method for mutational screen

*In silico* structure preparation and the Rosetta-Vienna RNP ΔΔG method was performed as previously described (31) but with the yeast UPF1-CH-HD structure (2XZL), and the results were compared to that of the UPF1-HD mutational screen (31). Chimera (36) was used to align human (2XZO) and yeast (2XZL) structures for residue conversions.

### Site-directed mutagenesis of UPF1-CH-HD plasmids for bacterial expression

100 ng of the pET28 vector harboring UPF1-CH-HD with N-terminal 6xHis and calmodulin binding peptide tags was used as the template. 500 nM of the appropriate primers designed using NEBaseChanger were added along with Phusion high-fidelity DNA polymerase, Phusion GC buffer, and 3% DMSO. PCR was initiated at 98°C for 5 min followed by 25 cycles of 98°C 1 min, T_a_ minus 5-6°C 30 sec, 72°C 5 min, and a final 72°C extension for 10 min. Primers are listed in **Supplementary Table S1**.

### Site-directed mutagenesis and cut-and-paste cloning of CLIP-UPF1 plasmids for mammalian expression at endogenous UPF1 levels

The pcDNA5-FRT-TO vector containing N-terminal CLIP-tagged full length UPF1 was used as the template for site-directed mutagenesis as previously described (31). Since these plasmids overexpress CLIP-UPF1 protein when stably expressed, the tetracycline-regulated promoter of pcDNA5-FRT-TO was replaced with the CMV promoter followed by a −40 kcal/mol hairpin 1 nt upstream of the predicted transcription start site using cut-and-paste cloning. Primers are listed in **Supplementary Table S1**.

### Mammalian stable cell line generation, protein extraction from cell lines, CLIP tag labeling, siRNA treatment, RNA extraction, cDNA synthesis, and RT-qPCR

These experiments were performed as previously described (31). RT-qPCR primers are listed in **Supplementary Table S2**.

### CLIP-UPF1 RNP extract preparation

Preparation was performed based on previous protocols (7, 37). Cells expressing CLIP-UPF1 mutants were pelleted at 1000 *g* for 3 min at 4°C, resuspended in 1 mL ice-cold 1X DPBS, then pelleted again at 2500 *g* for 1 min at 4°C. Excess supernatant was removed to avoid salt in subsequent next steps. Cell pellets were weighed on an analytical balance and resuspended in 5 cell volumes of ice-cold complete HLB (20 mM HEPES-NaOH pH 7.6, 2 mM MgCl_2_, 10% glycerol, 1 mM DTT, 1X Pierce Protease and Phosphatase inhibitor cocktail). Samples were incubated on ice for at least 5 min, subjected to two rounds of snap freezing in liquid N_2_ and thawing in a 37°C water bath, and placed on ice after the second round. NaCl was then added to a final concentration of 150 mM. Samples were incubated on ice for 5 min and centrifuged at 18,500 *g* for 15 min at 4°C. Supernatants containing intact RNA-protein complexes were transferred into a fresh cooled tube and were snap frozen in liquid N_2_. Protein concentration was determined via the Pierce 660 assay (Thermo Scientific).

### UPF2 immunoprecipitation (IP)

For each IP, 25 µL Protein G Dynabeads (Invitrogen) were washed three times with 500 µL bead wash buffer (20 mM HEPES-NaOH pH 7.6, 2 mM MgCl_2_, 10% glycerol, 150 mM NaCl, 0.1% NP-40) and resuspended in 25 µL bead wash buffer after the final wash. 10 µg of rabbit anti-RENT2 (Bethyl A303-929A) or rabbit IgG as a control for non-specific binding was mixed with the equilibrated beads and rotated end-over-end at room temperature for 1 hr to conjugate the beads to the antibody. The conjugated beads were then washed three times in 500 µL bead wash buffer and resuspended in 25 µL bead wash buffer. 5 mg/mL CLIP-UPF1 RNP extract (prepared as above) was added to the equilibrated conjugated beads, and the volume was brought up to 50 µL in bead wash buffer. Samples were rotated end-over-end at 4°C for 2 hr to bind endogenous UPF2 and washed three times in 500 µL bead wash buffer at 4°C. After the final wash, beads were resuspended in 10 µL of 1X NuPAGE LDS Sample Buffer (Invitrogen) and incubated at room temperature for 10 min. Supernatants (immunoprecipitated sample) were stored at −80°C for western blot analysis.

### Western blotting

Western blotting was performed as previously described (31). 1:1000 rabbit anti-RENT2 (Bethyl A303-929A) was used for UPF2 detection, 1:5000 goat anti-Rent1 (Bethyl A300-038A) was used for UPF1 detection, 1:1000 rabbit anti-phospho-(Ser/Thr) ATM/ATR substrate antibody (Cell Signaling 2851) was used for phospho-UPF1 detection, and 1:5000 mouse anti-beta actin (Cell Signaling 3700) was used for beta actin detection. Donkey anti-rabbit Alexa Fluor 680, donkey anti-goat AlexaFluor 680, or goat anti-mouse Alexa Fluor 680 secondary antibodies were diluted 1:10,000.

### rRNA depletion for RNA-seq libraries

Input total RNA was extracted as above using TRIzol Reagent followed by Shrimp dsDNase treatment and RNAclean XP bead cleanup and equilibrated to 100 ng/µL in nuclease-free water. rRNA depletion was adapted from previous work (38) and was performed in batches of 24 samples. An RNase H master mix containing 72 µL 5X RNase H buffer (250 mM Tris-HCl pH 7.5, 500 mM NaCl), 108 µL 100 mM MgCl_2_, and 108 µL nuclease-free water was pre-heated to 65°C. A 20 µL hybridization reaction containing 1X RNase H buffer (50 mM Tris-HCl pH 7.5, 100 mM NaCl), 50 µM EDTA, 5 µg homemade rRNA depletion oligonucleotide pool (see **Supplementary Table S3**), and 500 ng input RNA was subject to the following thermocycle to hybridize rRNA to the oligonucleotide pool: 95°C for 3 min, slow cool down of 0.1°C/sec to 65°C, 65°C for 5 min. During the 65°C 5 min step, 72 µL of either Hybridase Thermostable RNase H (Lucigen H39500) or Thermostable RNase H (NEB M0523) was added to the pre-heated RNase H master mix, thoroughly mixed, aliquoted 40 µL per tube in a pre-heated strip of 8 PCR tubes, and placed at 65°C. After the 65°C 5 min step, 10 µL RNase H master mix was added into each tube with a multichannel pipette and thoroughly mixed. Samples were then incubated at 65°C for 10 min to degrade rRNA and placed on ice. Samples were then cleaned up using 2.2X (66 µL) RNAClean XP beads (Beckman Coulter) that were equilibrated to 24°C for at least 30 min before use. Beads were mixed with samples, incubated 15 min at room temperature, then placed on an NEBNext Magnetic Separation Rack (NEB S1515S) for 5 min. On the magnetic rack, supernatant was removed and two washes of 200 µL freshly prepared 80% ethanol were performed without disturbing the bead pellet. Samples were briefly spun down, and residual ethanol was removed. Beads were then air dried for 4-5 min, resuspended with 41 µL nuclease-free water off the magnetic rack, and incubated for 5 min at room temperature. Tubes were then placed back on the magnetic rack. Once the solutions became clear, 37.5 µL of each supernatant was transferred into a fresh tube. The rRNA depletion oligonucleotides were then degraded by adding 7.5 µL of TURBO^TM^ DNase (2U/µL, Invitrogen AM2238) and 5 µL of the supplied 10X Reaction Buffer to each sample, mixed, incubated at 37°C for 30 min, and placed on ice. Samples were cleaned up as above but with 110 µL RNAClean XP beads and resuspended in 10 µL nuclease-free water after the ethanol washes, of which 6-7 µL was transferred to a fresh tube and stored at −80°C.

### RNA-seq library preparation

rRNA-depleted RNA was used as input and library preparation was performed using the xGen^TM^ RNA Library Preparation Kit (IDT 10009814) following the manufacturer’s instructions. Briefly, 1 µL Reagent F1, 2 µL Reagent F2, 4 µL Buffer F3, and 2 µL Reagent F4 were added to 5-6 µL rRNA-depleted RNA, and fragmentation was performed by incubating at 94°C for 10 min and immediately transferred to ice. Reverse transcription was performed by adding 1 µL Enzyme R1, 1 µL Enzyme R2, and 4 µL nuclease-free water to the fragmented samples, and subjected to the following thermocycle: 25°C 10 min, 42°C 30 min, 70°C 15 min, 4°C hold. Samples were then subjected to exonuclease treatment by adding 2.5 µL Exonuclease I (NEB M0293L) and 2.5 µL 10X Exonuclease I Reaction Buffer and incubating at 37°C for 30 min. Samples were equilibrated to 50 µL with the provided Low EDTA TE buffer and cleaned up using 1X (50 µL) SPRIselect beads (Beckman Coulter B23318) that were equilibrated to room temperature for at least 30 min before use. Beads were mixed with samples, incubated 5 min at room temperature, and placed on an NEBNext Magnetic Separation Rack (NEB S1515S) until the solution appeared clear. On the magnetic rack, supernatants were removed, and two washes of 200 µL freshly prepared 80% ethanol were performed without disturbing the bead pellet. Samples were briefly spun down, and residual ethanol was removed. Beads were then air dried for 3-5 min, resuspended with 12 µL Low EDTA TE off the magnetic rack, and incubated for 2 min at room temperature. Tubes were then placed back on the magnetic rack. Once the solutions became clear, 10 µL of each supernatant was transferred into a fresh tube, and samples were either stored at −20°C or subjected to the next step. Samples were denatured at 95°C for 2 min then immediately placed on ice for at least 2 min. Adaptase treatment was performed by adding 2 µL Buffer A1, 2 µL Reagent A2, 1.25 µL Reagent A3, 0.5 µL Enzyme A4, 0.5 µL Enzyme A5, and 4.25 µL Low EDTA TE to the denatured samples and thermocycled as follows: 37°C for 15 min, 95°C for 2 min, 4°C hold. 25 µL of the PCR Master Mix and 5 µL of a unique primer pair from either the xGen CDI Primers 96rxn (IDT 10009815) or xGen UDI 10 nt Primers (IDT 10008052) kits were added for indexing PCR, and the following thermocycle was performed: 1) 98°C 2 min, 2) 98°C 20 sec, 3) 60°C 30 sec, 4) 72°C 30 sec, repeat steps 2-4 13 times, 5) 72°C 1 min, 4°C hold. Samples were cleaned up as above but by adding 40.4 µL SPRIselect beads, resuspending in 22 µL Low EDTA TE, and transferring 20 µL to fresh tubes. Samples were quantified using the Quant-iT^TM^ PicoGreen dsDNA Assay Kit (Invitrogen P7589) following the manufacturer’s instructions, equilibrated to the same concentrations, and pooled together. The pooled library was quality checked on an Agilent TapeStation System by mixing 3 µL D1000 Sample Buffer (Agilent 5190-6502) with 1 µL pooled library and running in D1000 ScreenTape (Agilent 5067-5582). Libraries were sequenced on the Illumina NovaSeq 6000 or NovaSeq X+ platforms.

## Results

### The open state of UPF1 binds nucleic acids more slowly than the closed state

UPF1 is thought to predominantly reside in an enzymatically autoinhibited closed state and transition to an activated open state upon UPF2 binding. Enhancement of UPF1 catalytic activity by UPF2 is a conserved feature of nonsense-mediated mRNA decay (NMD) (21, 24, 39–41), but the functional consequences of UPF1 catalytic inhibition and activation remain poorly understood. Moreover, recent work has indicated that UPF2 also reduces UPF1 affinity for RNA (29). Therefore, we set out to systematically investigate the biochemical and functional effects of the structural rearrangement of UPF1 induced by UPF2 binding. To do so, we first kinetically probed how UPF1 interacts with nucleic acids in the closed and open states (**Figure 1A**). To study the open state of UPF1 without confounding factors such as binding and dissociation of UPF2, we generated mutants to disrupt the F192-I757 intramolecular interaction that normally stabilizes the closed state via docking of the CH domain onto the RecA2 domain (**Figure 1A**). We used the perpetually open F192E mutant (10, 24, 25) and a second mutant, I757R, predicted to disrupt this interaction (**Figure 1B**). We introduced these mutants into a well-characterized UPF1 construct containing the CH and helicase domains (UPF1-CH-HD; **Supplementary Figure S1A**, left).

**Figure 1.**
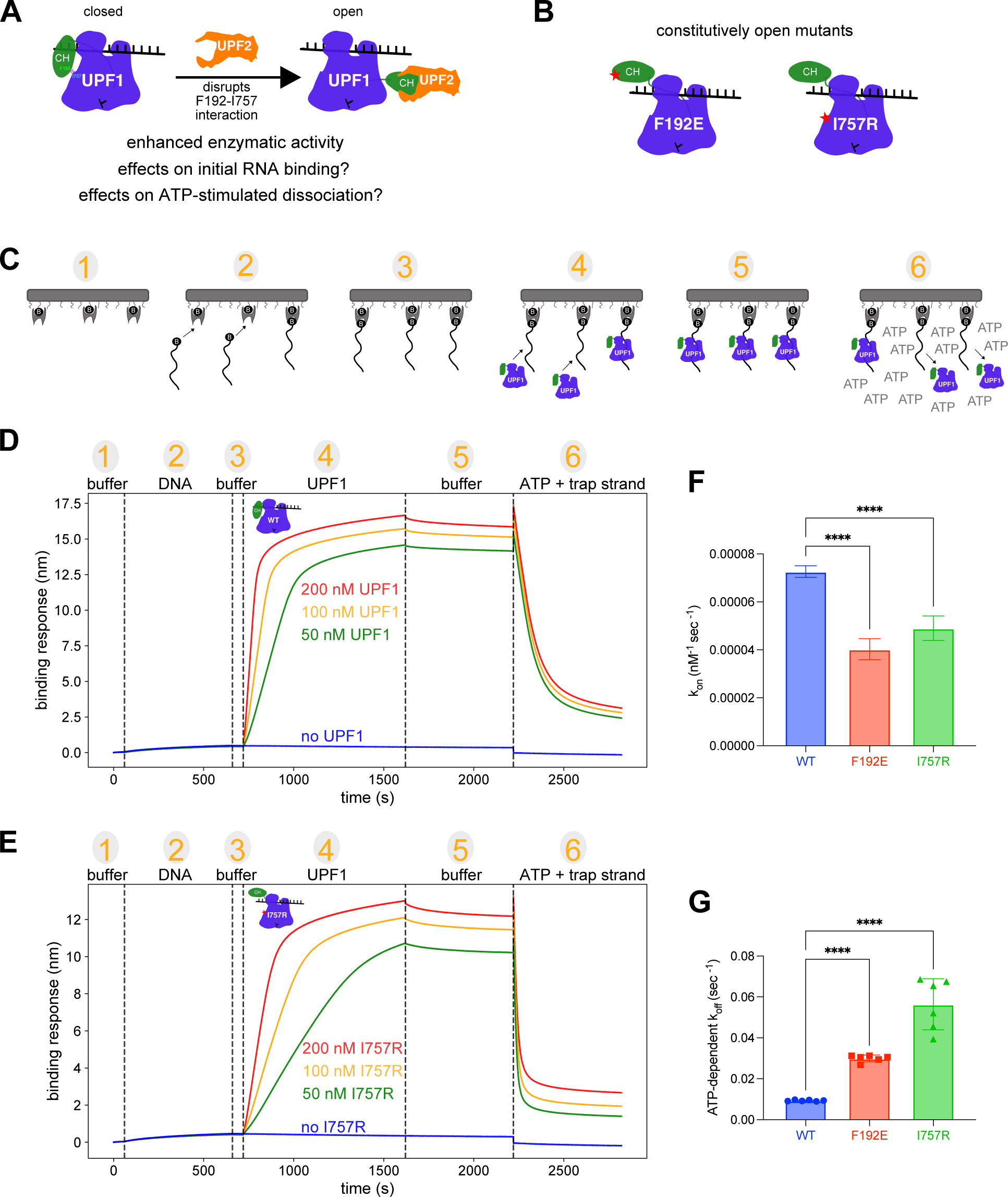
Open UPF1-CH-HD mutants exhibit slower nucleic acid binding but faster ATP-stimulated dissociation. (**A**) Schematic of possible biochemical and biophysical differences between the closed and UPF2-induced open states of UPF1. (**B**) Schematic of disinhibited open UPF1 mutants, F192E and I757R. (**C**) Schematic of the Octet Bio-Layer Interferometry (BLI) system, using biotin-streptavidin coated surface sensors (1). Sensors were sequentially dipped into wells with 10 nM of 50 nt biotinylated DNA oligonucleotide (2), buffer (3), varying concentrations of UPF1-CH-HD (4), buffer (5), and 2 mM ATP and 50 µM of 51 nt trap strand (not shown for clarity) (6). (**D-E**) Representative BLI traces of UPF1-CH-HD WT (**D**) and I757R (**E**). (**F-G**) Nucleic acid on-rates (**F**) and ATP-stimulated off-rates (**G**) calculated from BLI traces from 3 independent experiments. Error bars represent standard deviation. An ordinary one-way ANOVA statistical test was performed, with comparison between WT and each mutant (\*\*\*\**P* < 0.0001).

We first set out to quantitatively measure how the open state of UPF1 influences initial binding to nucleic acids utilizing the Octet Bio-Layer Interferometry (BLI) system. In this assay, binding of molecules to a streptavidin-coated sensor surface creates a wavelength shift proportional to the thickness of surface-bound molecules. We tethered a biotinylated DNA oligonucleotide substrate to the sensor, which resulted in a small thickness increase due to the small size of the oligonucleotide (**Figure 1C-D**, steps 1-3; **Supplementary Figure S1B-C**, steps 1-3). After addition of UPF1-CH-HD, the signal dramatically increased in a dose- and oligo-dependent manner (**Figure 1C-D**, step 4; **Supplementary Figure S1B-C**, step 4), indicating UPF1-CH-HD was specifically binding the substrate rather than non-specifically sticking to the surface. With a robust pre-steady state binding assay, we compared WT to the perpetually open mutants F192E and I757R (**Figure 1B**) and found the rates of F192E and I757R association with the substrate were significantly slower compared to WT (**Figure 1C-E**, step 4; **Supplementary Figure S1D**; **Figure 1F**). This effect was not influenced by nucleotide binding, as UPF1 association to the substrate was unchanged in the presence of the non-hydrolyzable ATP analog, AMP-PNP (**Supplementary Figure S1E-F**).

We corroborated the slow nucleic acid binding kinetics of the constitutively open mutants by adapting an established fluorescence-based unwinding assay. In this assay, displacement of a fluorescently labeled oligonucleotide strand by UPF1 results in quenching by a complementary oligonucleotide (**Supplementary Figure S2A**). With this assay, we first confirmed that a commonly used minimal UPF2 fragment (UPF2-S, containing the third MIF4G domain and UPF1 binding domain; **Supplementary Figure S1A**, right) enhanced UPF1-CH-HD unwinding on various DNA and RNA substrates in a dose-dependent manner (**Supplementary Figure S2B-C**) but did not significantly affect UPF1-CH-HD relative processivity (**Supplementary Figure S2D-E**, **Supplementary Materials & Methods**). However, we observed drastically different unwinding kinetics when we varied the duration of incubation of UPF1 or UPF1-UPF2 complexes with the unwinding assay substrate (**Supplementary Figure S2F**). Unwinding activity increased markedly with increasing incubation time for UPF1-CH-HD/UPF2-S complexes but not UPF1-CH-HD alone (**Supplementary Figure S2G-I**), indicating that the complex may bind the substrate more slowly. To distinguish between effects caused by UPF2 binding and those caused by the open state of UPF1, we mimicked the open state by using a UPF1 construct lacking the CH domain (UPF1-HD; **Supplementary Figure S1A**, left). Like UPF1-UPF2 complexes, UPF1-HD showed enhanced sensitivity to incubation time for unwinding activity (**Supplementary Figure S2I**), suggesting that slow nucleic acid binding kinetics are an intrinsic property of the open state of UPF1. This result was further supported by fluorescence anisotropy equilibrium binding assays, in which we observed reduced nucleic acid binding affinity of F192E compared to WT UPF1-CH-HD in both the presence and absence of ATP (**Supplementary Figure S2J**).

### UPF2 enhances UPF1 ATP-stimulated dissociation from nucleic acids by promoting the open state

The open state induced by UPF2 appears to delay initial UPF1 binding to nucleic acids (**Figure 1F**) but also enhances UPF1 enzymatic activity on nucleic acids (**Supplementary Figure S2B-C**) (21, 24). We next investigated how UPF2 affects UPF1 dissociation from nucleic acids, since UPF1 dissociation from RNA is thought to be an important determinant of NMD efficiency and specificity (5, 30–32, 35, 42). Using BLI, we incubated the sensors containing UPF1-bound DNA in buffer (**Figure 1C**, step 5), which caused negligible ATP-independent dissociation of UPF1 (**Figure 1D-E**, step 5). When we moved the complex into a solution with saturating ATP and trap strand to capture any dissociated UPF1 molecules (**Figure 1C**, step 6), we observed rapid ATP-stimulated dissociation (**Figure 1D**, step 6), which was significantly faster with the open F192E and I757R mutants (**Figure 1E**, step 6; **Figure 1G**).

To corroborate the finding that constitutively open mutants undergo accelerated ATP-stimulated dissociation, we used a previously developed fluorescence anisotropy-based dissociation (FAD) assay. In this assay, fluorescence polarization decreases over time as UPF1 molecules dissociate from the fluorescent substrate and are captured by the excess trap strand in solution (**Figure 2A**) (31). With this assay, we found that UPF1-CH-HD underwent efficient ATP-stimulated dissociation from the substrate (**Figure 2B**, orange curve), and UPF2-S enhanced this dissociation (**Figure 2B**, red curve). We also performed control experiments to verify that UPF2-S did not detectably bind the fluorescent substrate under these assay conditions (**Supplementary Figure S3A**). Additionally, we observed high rates of UPF2-independent ATP-stimulated dissociation of I757R and F192E (**Figure 2C-D**). These findings were recapitulated with structured substrates (**Supplementary Figure S3B**) and at a higher pH (**Supplementary Figure S3C**).

**Figure 2.**
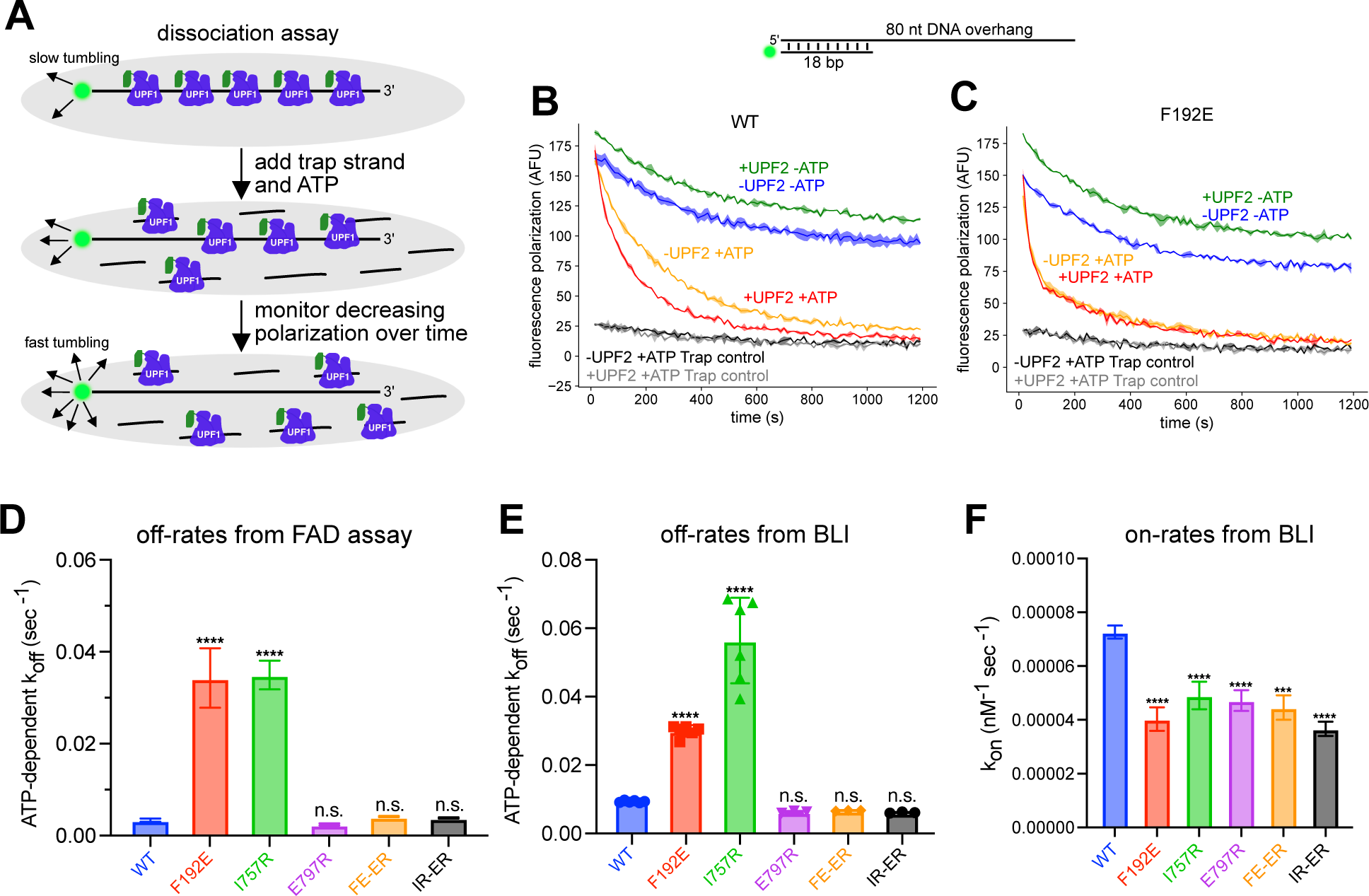
UPF1-CH-HD exhibits efficient ATP-stimulated dissociation from nucleic acids, an activity enhanced in the open state. (**A**) Schematic of the fluorescence anisotropy-based dissociation (FAD) assay. UPF1-CH-HD was in 5-fold molar excess (71.25 nM) over the substrate (14.25 nM) and UPF2-S (not shown) was in 10-fold molar excess (142.5 nM) over the substrate unless otherwise indicated. Trap strand was added at 1000-fold molar excess (14.25 µM) to the substrate immediately prior to injection of ATP. (**B-C**) Representative curves of FAD of WT UPF1-CH-HD (**B**) or F192E (**C**) with an 80 nt 3′ DNA overhang substrate. 1000-fold molar excess of trap strand was added before UPF1-CH-HD addition to the substrate in trap controls (black and gray curves). (**D**) ATP-dependent off-rates calculated from FAD assays of different mutants from 2 independent experiments. Error bars represent standard deviation. An ordinary one-way ANOVA statistical test was performed, with comparison between WT and each mutant (\*\*\*\**P* < 0.0001). (**E-F**) As in Figure 1G (**E**) and Figure 1F (**F**), from BLI experiments with the indicated mutants from 2 independent experiments. Error bars represent standard deviation. An ordinary one-way ANOVA statistical test was performed, with comparison between WT and each mutant (\*\*\**P* < 0.001, \*\*\*\**P* < 0.0001).

The higher rates of ATP-stimulated dissociation of F192E and I757R in the absence of UPF2-S (**Figure 2D**) suggest that UPF2 primarily promotes UPF1 dissociation from nucleic acids by stimulating ATP hydrolysis (5, 18, 24, 39–41). We therefore predicted that slowing the ATP hydrolysis rates of the F192E and I757R proteins would correspondingly slow their dissociation. To test this hypothesis, we combined F192E or I757R with a mutation that decreases UPF1 ATPase rate, E797R (31). As predicted, the compound F192E/E797R (FE-ER) and I757R/E797R (IR-ER) mutants both underwent slower ATP-stimulated dissociation than the F192E and I757R mutants alone, with rates similar to WT in FAD (**Figure 2D**) and BLI assays (**Figure 2E**). The addition of the E797R mutation to the F192E and I757R mutant backgrounds did not alter nucleic acid on-rates (**Figure 2F**).

### In silico modeling predicts cellular UPF1-RNA interactions are dominated by intrinsic kinetic properties of UPF1 rather than UPF2-induced kinetic properties

UPF2 has the potential to both positively and negatively regulate UPF1 function in NMD by slowing down initial RNA binding, enhancing unwinding/translocation, and/or enhancing ATP-stimulated dissociation from RNA. However, these effects are likely constrained by the relative abundances of UPF1 and UPF2 in human cells, as proteomic studies estimate that UPF1 is approximately 10-fold more abundant than UPF2 in HEK-293T cells (∼450,000 versus ∼54,000 molecules per cell) (43, 44). To better understand the relative contributions of UPF2 to UPF1 occupancy on mRNAs in cells, we used our experimentally determined kinetic parameters of UPF1-RNA (**Figure 1F-G**; **Figure 2D-F**) and UPF1-UPF2 interactions (**Supplementary Figure S4A-D**) to simulate interactions among equimolar amounts of RNA and UPF1 and differing quantities of UPF2 (**Figure 3A**, **Supplementary Table S4**). Computational modeling predicted that UPF2 can only suppress the fraction of UPF1 bound to RNA at high UPF2:UPF1 stoichiometries **(Figure 3B**). Correspondingly, a large majority of UPF1 association (**Figure 3C**) and dissociation (**Figure 3D**) events are predicted to occur independent of UPF2 at the physiological UPF2:UPF1:mRNA stoichiometry of 1:10:10 (**Figure 3C-D**, 0.1 column). Taken together, intrinsic UPF1 properties are likely to be largely responsible for cellular RNA interactions, with minimal effects from UPF2 due to its lower relative abundance to UPF1.

**Figure 3.**
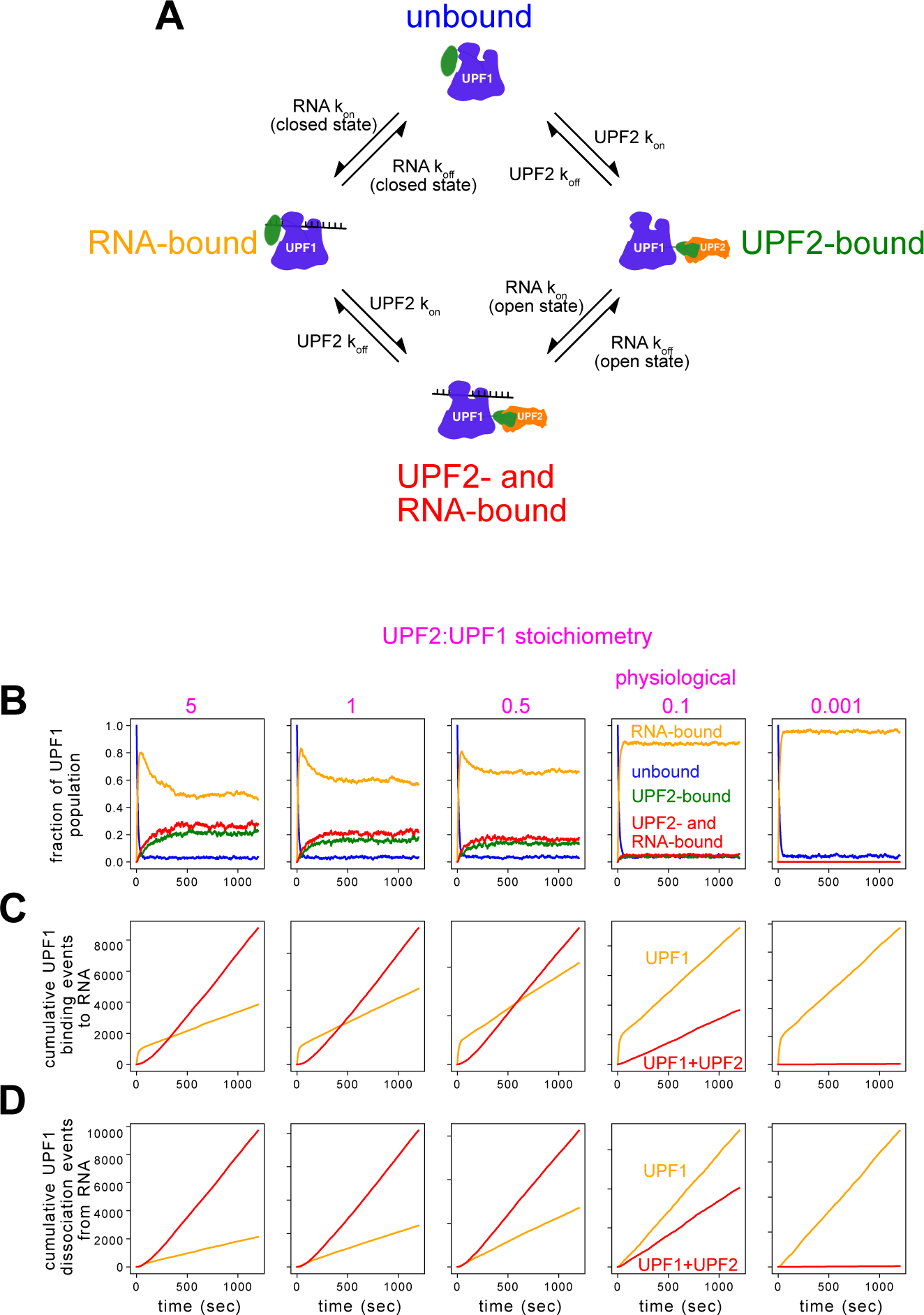
In silico modeling of UPF1-UPF2-RNA interactions. (**A**) Schematic of possible states of UPF1 in the simulations. Parameters used for each kinetic arrow are derived from BLI and FAD assays in the presence of saturating ATP (see **Supplementary Table S4**). (**B-D**) Simulation of 1000 UPF1s, 1000 RNAs, and differing amounts of UPF2s from 100,000 (left panels) to 1 (right panels), assuming saturating ATP conditions. The physiological UPF2:UPF1 stoichiometry is indicated, corresponding to simulations with 100 UPF2s. The state of UPF1 over time was monitored in simulations (**B**). Cumulative simulated RNA binding events (**C**) and dissociation events (**D**) of UPF1 alone (orange) or UPF2-bound UPF1 (red) plotted over time.

Our finding of efficient ATPase-stimulated release of UPF1 from nucleic acid in the absence of UPF2 (**Figure 1D**; **Figure 2B**) raised the question of whether this requires spontaneous undocking of the CH domain into the open state. Alternatively, UPF1 ATP hydrolysis and RNA release could occur at slower rates in the closed state compared to the open state. To gain insight into this question, we combined multiple turnover unwinding assays on different structured substrates (**Supplementary Figure S5A-C**) with computational kinetic modeling. We observed that UPF1 unwound short hairpin DNA (hpDNA) substrates with biphasic kinetics in multiple- and single-turnover conditions (**Supplementary Figure S5B**, **D**, **E**). We did not observe these biphasic kinetics with the open F192E mutant (**Supplementary Figure S5F**) or UPF1-HD (31). Computational modeling (**Supplementary Materials & Methods**) predicted that this kinetic profile is consistent with UPF1 unwinding 0.75 bp/sec in the open state and 0.025-0.1 bp/sec in the closed state, with a 0.005-0.1 probability of opening per step and a 0.0-0.5 probability of closing per step (**Supplementary Figure S5G**, panels boxed in red indicating at least one simulated curve matches the experimental data). This indicates that not only can UPF1 unwind without UPF2, but also has the ability to stochastically switch between the open and closed states, with activity in both states.

### Identification of more strongly autoinhibited UPF1 mutants

Based on the predicted central importance of intrinsic UPF1 properties in determining UPF1-RNA interaction dynamics (**Figure 3**), we pursued a strategy of identifying and/or characterizing UPF1 mutants with alterations in autoinhibition, ATP hydrolysis, and ATP-stimulated dissociation. This work was bolstered by our unexpected identification of a UPF1 residue located at the 3′ end of the RNA binding channel, G619 (**Figure 4A**), that can be mutated to generate proteins that are more strongly autoinhibited and are thus more reliant on UPF2 for enzymatic activation. In the course of characterizing hits from two parallel *in silico* screens (**Supplementary Figure S6A-C**; **Supplementary Table S5-6**) (31) to identify UPF1 mutants with differential RNA binding properties in the open and closed states, we found that mutation of G619 to histidine, phenylalanine, or tyrosine resulted in stronger autoinhibition of UPF1.

**Figure 4.**
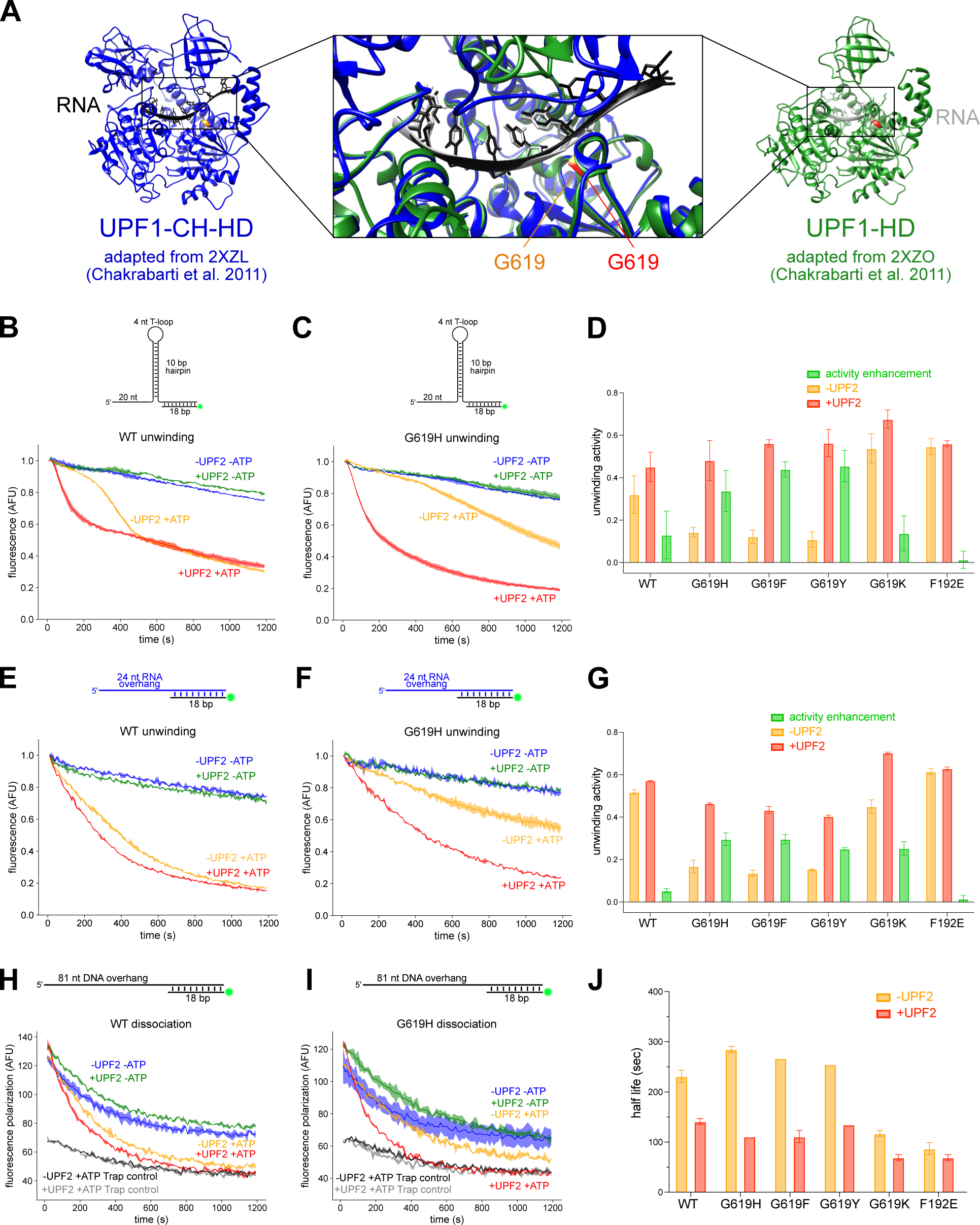
Mutating UPF1-CH-HD G619 to an aromatic amino acid increases autoinhibition. (**A**) Structures of UPF1-CH-HD (left, adapted from PDB 2XZL) and UPF1-HD (right, adapted from PDB 2XZO), following addition of missing residues and relaxation. The G619 residue is shown in orange for UPF1-CH-HD and red for UPF1-HD. A zoomed-in view of the structural context of G619 (middle). (**B-C**) Representative curves from multiple turnover unwinding assays using a short hpDNA substrate of UPF1-CH-HD WT (**B**) or G619H (**C**). (**D**) Quantification of unwinding activity from 3 independent experiments (see **Supplementary Materials & Methods**). Activity enhancement (green bars) was calculated by subtracting activity in the absence of UPF2 (orange bars) from activity in the presence of UPF2 (red bars). Error bars represent standard deviation. (**E-F**) As in (**B-C**) but with a 5′ RNA overhang substrate and with 4-fold molar excess of UPF2-S (330 nM). (**G**) As in (**D**) but quantified from unwinding experiments with a 5′ RNA overhang substrate from 2 independent experiments. (**H-I**) Representative curves of FAD with a 5′ DNA overhang substrate of UPF1-CH-HD WT (**H**) or G619H (**I**). 1000-fold molar excess of trap strand was added before UPF1-CH-HD addition to the substrate in trap controls (black and gray curves). (**J**) Calculation of half life for FAD assays of UPF1-CH-HD alone (orange) or in the presence of UPF2-S (red) from 2 independent experiments.

In the presence of UPF2-S, the G619H, G619F, and G619Y mutants exhibited unwinding activity comparable to WT UPF1-CH-HD on DNA (**Figure 4B-C**, **Supplementary Figure S7A-B**, red curves; **Figure 4D**, red bars) and RNA (**Figure 4E-F**, **Supplementary Figure S7C-D**, red curves; **Figure 4G**, red bars), but without UPF2-S, they showed significantly lower unwinding activity compared to WT in the absence of UPF2-S (**Figure 4B-G**, **Supplementary Figure S7A-D**, orange curves/bars). G619H, G619F, and G619Y mutants had minimal dissociation defects on DNA substrates with a 3′ overhang compared to WT (**Supplementary Figure S8A-E**) but exhibited modestly slower ATP-stimulated dissociation from substrates containing a double-stranded region (**Figure 4H-I**, **Supplementary Figure S7E-F**, compare orange curves to blue curves; **Figure 4J**, orange bars). This defect was overcome when UPF2-S was present (**Figure 4H-I**, **Supplementary Figure S7E-F**, compare red curves to green curves; **Figure 4J**, red bars). These mutants did not exhibit nucleic acid binding defects, as measured by fluorescence anisotropy (**Supplementary Figure S8F-I**) or increased Alexa Fluor 647-labeled UPF2-S binding, as measured by microscale thermophoresis (MST, **Supplementary Figure S8J**). Together, these findings suggest that the G619H, G619F, and G619Y mutants are more autoinhibited but that UPF2 binding is sufficient to overcome this enhanced autoinhibition.

Based on the positioning of residue 619 at the 3′ end of the RNA binding channel (**Figure 4A**), we speculated that enhanced autoinhibition may involve base stacking interactions between the nucleic acid substrate and the aromatic residue at position 619. In support of this, mutation to non-aromatic residues (i.e. G619K and G619C) were not more autoinhibited than WT in either unwinding (**Supplementary Figure S9A-F**) or dissociation assays (**Supplementary Figure S9G-J**). In fact, G619K had slightly higher unwinding activity (**Supplementary Figure S9A-C**) as previously reported for G619K in the UPF1-HD construct (31). Furthermore, mutating G619 did not appear to influence nucleic acid on-rate, as G619K and G619H both exhibited WT-like on-rates (**Supplementary Figure S9K-L**). Only G619K modestly enhanced ATP-dependent off-rates on single-stranded substrates in BLI experiments (**Supplementary Figure S9M**).

### More strongly autoinhibited UPF1 mutants support regulation of canonical NMD targets

To test the effects of UPF1 autoinhibition and ATPase activation on cellular NMD, we assembled a panel of new and previously characterized UPF1 mutants with alterations in various aspects of catalytic activation and ATP-stimulated dissociation. The panel comprises UPF1 mutants with 1) impaired ATP-stimulated dissociation (ATPase-deficient mutants K498A (19, 45), DE636AA (20, 46), and R843C (47, 48); mechanically impaired A546H (31, 49)), 2) impaired mechanical properties but restored ATP-stimulated dissociation (G619K/A546H [GA-KH] (31)), 3) increased autoinhibition (G619H, G619F, G619Y), 4) decreased autoinhibition with slower nucleic acid on-rates and accelerated ATP-stimulated dissociation (F192E, I757R), and 5) decreased autoinhibition with slower nucleic acid on-rates but WT-like rates of ATP-stimulated dissociation (FE-ER and IR-ER) (**Figure 5A**).

**Figure 5.**
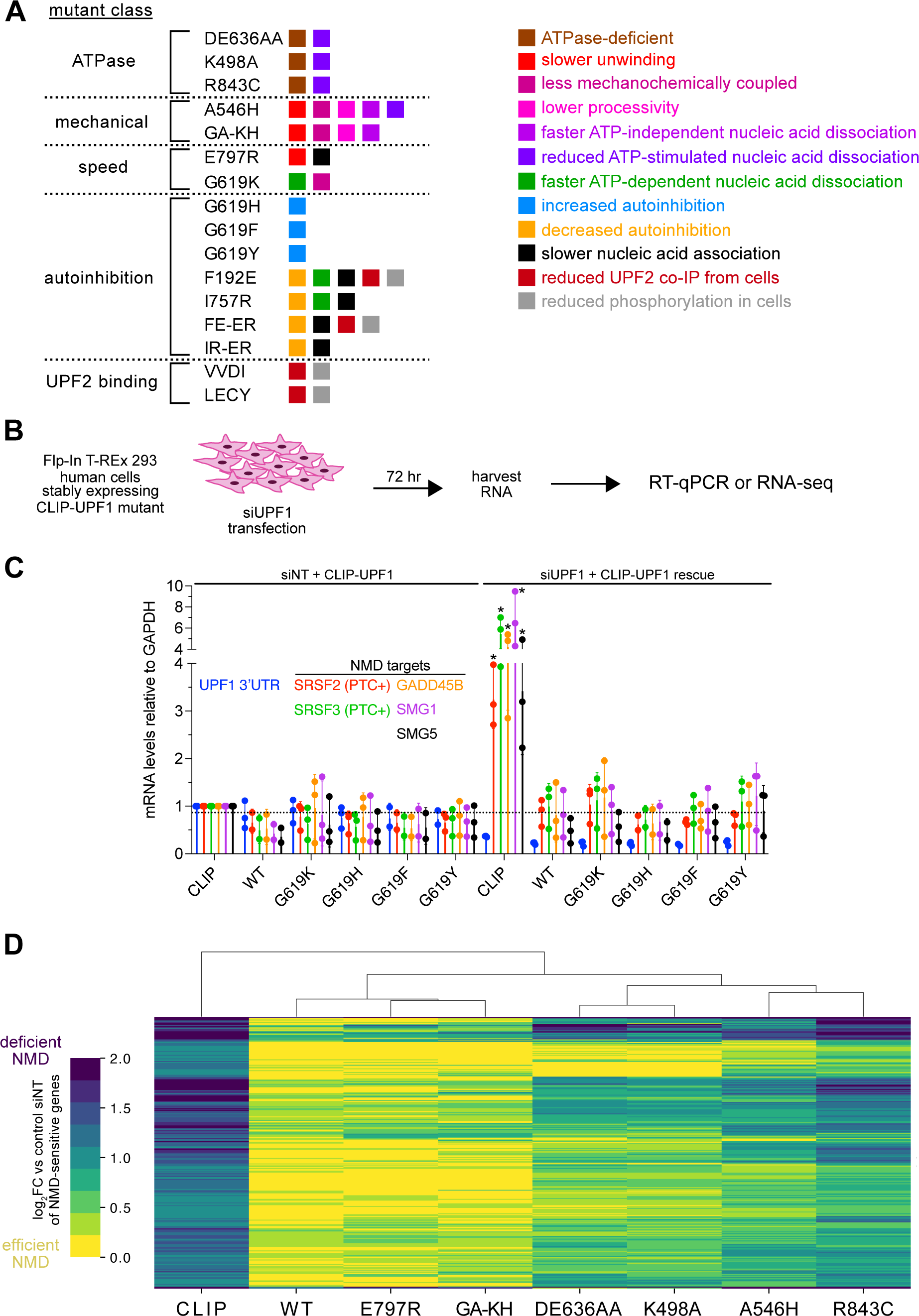
Impaired ATP-stimulated dissociation, but not increased autoinhibition of UPF1, perturbs mammalian NMD. (**A**) Qualitative classification of UPF1 mutants based on *in vitro* characterization from this study and previous work (31). Omission of a property does not mean the mutant does not have that property, as not all assays were performed using all mutants. (**B**) Schematic of knockdown/rescue experiments in Flp-In T-Rex 293 cells stably expressing CLIP-UPF1 mutants. Rescue efficiency was determined using RT-qPCR of canonical NMD target mRNAs or transcriptome-wide using RNA-seq. (**C**) mRNA levels of NMD targets relative to GAPDH using RT-qPCR normalized to the siNT CLIP condition from 3 independent experiments. UPF1 3′ UTR mRNA levels were measured to determine the extent of endogenous UPF1 knockdown. SRSF2 and SRSF3 primers were specific for PTC-containing exons. Error bars represent standard deviation. siNT indicates non-targeting siRNA and siUPF1 indicates UPF1 siRNA. A two-way ANOVA statistical test was performed, with comparison between each siUPF1 condition and the siNT CLIP condition (\**P* < 0.0001, **Supplementary Table S7**). (**D**) Gene-level analysis of RNA-seq data of knockdown/rescue experiments from 3 independent experiments, filtered for genes that were upregulated > 2-fold in the siUPF1 CLIP condition and rescued > 75% in the siUPF1 WT condition. All gene abundances were compared to the siNT CLIP control condition. Only endogenous UPF1 knockdown conditions are shown. Both rows (genes) and columns (mutants) were clustered using the default parameters of the seaborn clustermap python package (52). Row dendrogram is not shown for clarity. See **Supplementary Table S8** for raw data.

To evaluate cellular NMD functionality, we generated cell lines stably expressing N-terminal CLIP-tagged versions of these UPF1 mutants at levels comparable to endogenous UPF1 (31) and performed knockdown/rescue experiments followed by RT-qPCR or RNA-seq (**Figure 5B**). Knockdown of endogenous UPF1 caused up-regulation of several well-characterized NMD target mRNAs (e.g. containing a premature termination codon [PTC] or long 3′ UTR), an effect completely counteracted by expression of WT CLIP-UPF1 but not the CLIP tag alone (**Figure 5C**). Despite having a distinctly greater dependence on UPF2 for enzymatic activation *in vitro*, the more strongly autoinhibited G619H, G619F, and G619Y mutants fully rescued the effects of endogenous UPF1 knockdown (**Figure 5C**). Because these mutants were indistinguishable from WT UPF1 or the control G619K mutant in knockdown/rescue experiments, we chose to focus further tests of the butterfly model on UPF1 mutants with altered ATP-dependent dissociation from RNA.

### UPF1 mutants with impaired ATP-stimulated dissociation exhibit cellular NMD defects

In RNA-seq of knockdown/rescue experiments, we confirmed that UPF1 mutants with RNA dissociation defects did not restore NMD as well as WT UPF1. We evaluated UPF1 activity in two ways. First, we monitored overall changes in RNA abundance by gene-level differential expression analyses (**Figure 5D and Supplementary Figure S10A-C**). Second, we investigated isoform-specific effects of UPF1 knockdown and rescue by performing splice junction-level analyses of differential transcript isoform usage (**Supplementary Figure S10D-E**). This approach focuses on reads arising from cassette exon splicing events and thus enriches for transcriptome alterations linked to PTCs. NMD-deficient mutants in this class consist of A546H, which is defective for ATP-stimulated dissociation from structured substrates (31), and extensively characterized ATPase-deficient mutants K498A, DE636AA, and R843C. In contrast, E797R, which hydrolyzes ATP slowly, and the GA-KH compound mutant that exhibits A546H-like mechanical deficiencies but restored ATP-stimulated dissociation relative to A546H, promoted NMD, with similar efficiency as WT CLIP-UPF1 (**Figure 5D**, **Supplementary Figure S10A-E**). Together these results support the idea that cellular NMD relies on proper ATP-stimulated dissociation of UPF1.

### Constitutively open UPF1 mutant F192E impairs UPF2 association and phosphorylation

We next set out to determine the functional consequences of constitutive relief of UPF1 autoinhibition. As part of our initial cellular characterization of UPF1 mutants, we assayed interactions between selected representatives of the UPF1 mutant classes and UPF2 by immunoprecipitation of endogenous UPF2. Along with selected mutants from above, we included two additional UPF1 mutants as positive controls, VV204-5DI (VVDI) and LECY181-4VRVD (LECY), previously found to have moderate and severe reductions in UPF2 binding, respectively (13). Neither VVDI nor LECY exhibited UPF2 binding within the detection limit of this assay (**Supplementary Figure S11A**), and both correspondingly showed a drastic reduction in steady-state phosphorylation (**Supplementary Figure S11B-C**). I757R, IR-ER, E797R, and G619H retained full association with UPF2, but F192E and FE-ER exhibited reduced co-immunoprecipitation with UPF2 (**Supplementary Figure S11A**,**E**), consistent with evidence of direct binding of F192 by UPF2 (25). F192E and FE-ER were correspondingly phosphorylated at significantly lower levels, likely due to their reduced association with UPF2 (**Supplementary Figure S11F-G**).

### Constitutively open UPF1 mutants are partially defective for cellular NMD

In the RNA-seq of knockdown/rescue experiments, I757R restored NMD to a greater extent than F192E, as determined by gene-level differential expression analyses (**Figure 6A-B**) and splice junction-level analyses of NMD-sensitive transcript isoforms (**Figure 6C**). Lower NMD activity of F192E is consistent with our observation of impaired F192E UPF2 association and phosphorylation (**Supplementary Figure S11A-G**). F192E was less active than the WT and E797R positive controls on well-established NMD targets but more active than LECY (**Supplementary Figure S11H**), suggesting that F192E retains some ability to recruit UPF2 (**Figure 6A-C**, **Supplementary Figure S11H**). This is consistent with impaired UPF2 binding and phosphorylation of VVDI (**Supplementary Figure S11A-C**), yet moderate restoration of NMD (**Supplementary Figure S11H**). The compound FE-ER mutant more efficiently restored NMD transcriptome-wide relative to F192E, such that it attained similar activity to I757R (**Figure 6A-C**). Addition of the E797R mutation in the I757R background (i.e. IR-ER) yielded no further increase in NMD rescue compared to I757R (**Figure 6A-C**), indicating that the constitutively activated mutants have defects that cannot be overcome by slowing the rate of ATP hydrolysis. Together, our data suggest that constitutive undocking of the CH domain has multiple impacts on UPF1 biochemical activities but can be partially alleviated by reducing the rate of ATP-stimulated dissociation to WT-like levels.

**Figure 6.**
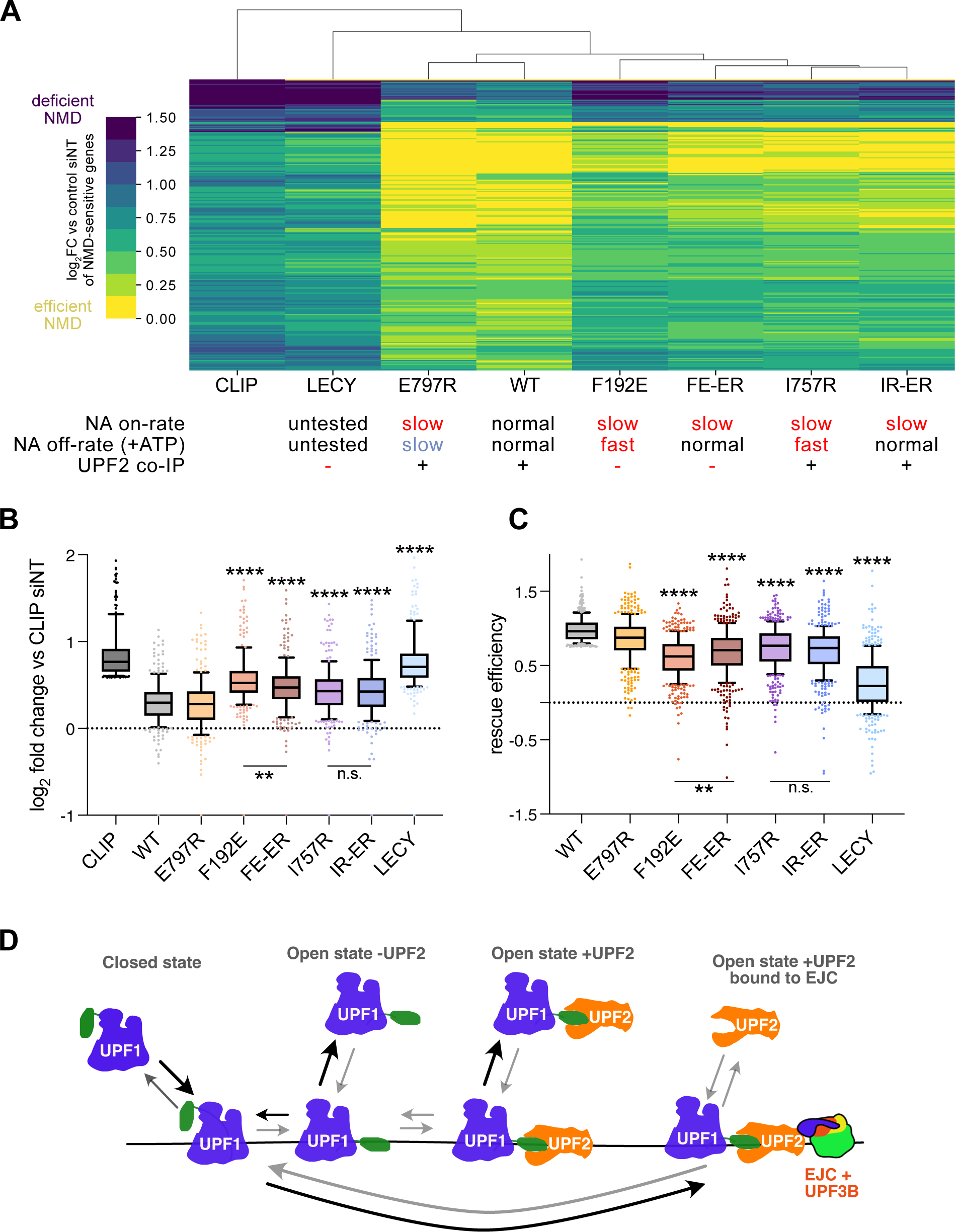
Constitutively open UPF1 mutants impair cellular NMD. (**A**) Data acquisition and analysis performed as in Figure 5D (minor differences described in **Supplementary Materials & Methods**), with the indicated set of UPF1 mutants assayed in 3 independent experiments. Only genes that were upregulated > 1.5-fold in siUPF1 CLIP and rescued > 50% in siUPF1 WT are shown. Only endogenous UPF1 knockdown conditions are shown. Under each mutant are qualitative categorizations of nucleic acid (NA) binding kinetics and ability to co-immunoprecipitate (co-IP) UPF2 from cell extracts. The LECY mutant was not tested *in vitro*. (**B**) As in (**A**) but visualized on a boxplot, with boxes indicating interquartile ranges and whiskers indicating 10-90% ranges. Kruskal-Wallis tests with Dunn’s correction for multiple comparisons were performed between E797R and other conditions, F192E and FE-ER, and I757R and IR-ER. CLIP and WT were excluded from statistical analysis as their distributions were truncated due to filtering (\*\*\*\**FDR* < 0.0001, *\*\*FDR* < 0.01). (**C**) rMATS analysis of RNA-seq data to analyze exon inclusion events. Only transcripts that increased in inclusion by 10% in siUPF1 CLIP and underwent > 75% rescue upon expression of WT UPF1 are shown. Boxes indicate interquartile ranges and whiskers indicate 10-90% ranges. Kruskal-Wallis tests with Dunn’s correction for multiple comparisons were performed between E797R and other conditions, F192E and FE-ER, and I757R and IR-ER. CLIP and WT conditions were excluded from statistical analysis as their distributions were truncated due to filtering (\*\*\*\**FDR* < 0.0001, *\*\*FDR* < 0.01). See **Supplementary Table S8** for raw data. (**D**) Proposed model of the effects of UPF1 conformational changes on binding kinetics in NMD. Arrow weights are relative within each UPF1 state based on BLI and FAD kinetic parameters. Bottom arrows between UPF1 closed state -UPF2 and UPF1 open state +UPF2 are not based on kinetic experiments but reflect expected enhancement of UPF1-UPF2 interactions through UPF2 tethering to the EJC.

## Discussion

The fact that autoinhibition of UPF1 catalytic activity is conserved throughout eukaryotes implies that it plays an important role in cellular NMD (24). However, the mechanisms by which UPF1 autoinhibition and activation by UPF2 promote mRNA decay are poorly understood. To address this gap, we used a combination of biochemical/biophysical, computational, and sequencing-based methods to identify and characterize UPF1 mutants. We find that the open state of UPF1 has slower association kinetics and faster ATP-stimulated dissociation kinetics to and from nucleic acids, respectively (**Figures 1, 2 and 6D**). Computational modeling based on kinetic measurements suggest that the low cellular abundance of UPF2 limits its impact on UPF1-RNA interactions. At cellular UPF2:UPF1:mRNA stoichiometries, modeling predicts that the majority of UPF1-RNA binding and dissociation events occur in the absence of UPF2 (**Figure 3**). Nonetheless, we find that newly identified UPF1 mutants that are more strongly inhibited in the absence of UPF2 (**Figure 4**) are able to support decay of canonical NMD target mRNAs (**Figure 5**). In contrast, UPF1 mutants that are unable to undergo efficient ATP-stimulated dissociation or are perpetually in the open state have reduced NMD activity in human cells (**Figures 5 and 6**). These findings are consistent with the recently proposed butterfly model of NMD, in which an important function of UPF1 ATPase activity is autoregulation of RNA binding (31).

Importantly, UPF1 does not require UPF2 for ATP hydrolysis or ATP-stimulated dissociation. Instead, UPF2 accelerates the rates of both processes (**Figure 2 and Figure 6D**) (5, 18, 24, 39–41). This implies that UPF1 sampling of the transcriptome through successive rounds of binding and dissociation need not require activation by UPF2. In line with this, modeling of interactions among UPF1, UPF2, and RNA that takes into account experimentally derived kinetic parameters and cellular abundance of these components predicts that a large fraction of UPF1-RNA association and dissociation events are UPF2-independent (**Figure 3**). Further, we leverage the kinetic measurements of *in vitro* hairpin unwinding experiments to perform modeling that indicates UPF1 catalytic activity can occur in both the open and closed states (**Supplementary Figure S5**). These results blur the line between the previously proposed RNA-clamping versus RNA-unwinding modes of the closed versus open states, respectively (24). More in-depth experiments to directly interrogate the relationship between CH domain undocking and enzymatic activity will be required to further elucidate opening and closing dynamics.

While our experimental data and computational modeling suggest that most cellular UPF1-RNA association and dissociation events are UPF2-independent, the inhibitory effects of UPF2 binding on UPF1 may still be important in multiple ways. For instance, slow UPF1-UPF2 association with RNA may help to impose an order of assembly such that UPF1 binding to potential substrates is followed by UPF2 binding. This mechanism may help prevent promiscuous RNA decay, as it disfavors initial binding between mRNAs and the pro-decay UPF1-UPF2 complex. This mechanism could also enforce preferential decay of mRNAs harboring one or more exon junction complexes (EJCs), as UPF2 recruitment to EJCs would in turn drive UPF1-UPF2 interactions, UPF1 phosphorylation, and decay. A second mechanism by which UPF2 may help enforce decay specificity is by stimulating dissociation of UPF1. UPF2 association with UPF1 bound in a context that does not meet additional requirements for decay may help to recycle UPF1 off RNAs more quickly. In this scenario, UPF2 would serve a proofreading role to prevent inappropriate decay by decreasing the kinetic window for decay initiation.

The data presented here also reinforce previous findings that UPF1 mutants with impaired ability to perform work in the form of unwinding and protein displacement can be fully competent for NMD (31). The GA-KH double mutant unwinds slowly (∼0.25 bp/sec) with poor processivity (∼10-20 bp per binding event) and is mechanochemically uncoupled (31), yet almost precisely phenocopies wild-type UPF1 transcriptome-wide in knockdown/rescue experiments (**Figure 5**). Likewise, the E797R mutation slows the rates of ATP hydrolysis and translocation (∼0.2 bp/sec) but permits equal or modestly greater NMD rescue following endogenous UPF1 knockdown (**Figures 5 and 6**). These findings are compatible with ATP-dependent roles for UPF1 in local RNP remodeling or ribosome recycling (22, 23), but do strongly argue against models in which UPF1 acts as a processive RNP remodeler to promote RNA degradation. Further, the substantial activity of constitutively active mutants in cells (**Figure 6**), suggest that specific activation of UPF1 enzymatic activity in response to UPF2 binding or other triggers is largely dispensable for NMD.

The partial improvement in NMD activity of the FE-ER double-mutant relative to the F192E mutant alone is consistent with the hypothesis that one function of UPF1 autoinhibition is to slow the rate of ATP-stimulated dissociation. However, we did not see a similar effect for the IR-ER double-mutant versus I757R (**Figure 6**). Given the multifaceted roles of UPF1 in NMD, it is likely that constitutively open UPF1 mutants have impaired activity for multiple reasons, some of which can be ameliorated by depressing overall ATPase activity and some of which cannot.

Interestingly, both I757R and IR-ER both showed similar NMD efficiencies as FE-ER, suggesting that WT levels of UPF2 binding and phosphorylation may insulate them from other defects. We identified four relevant properties of F192E that may contribute to decreased activity in NMD: slower nucleic acid on-rate, faster nucleic acid ATP-stimulated off-rate, reduced UPF2 association, and reduced phosphorylation. I757R and IR-ER were stably associated with UPF2 in co-immunoprecipitation experiments and were phosphorylated at levels comparable to WT UPF1. Like F192E and FE-ER, the on-rates of I757R and IR-ER were reduced, and we cannot exclude the possibility that other aspects of the mutant proteins’ interactions, folding, or enzymatic activities are perturbed. Future investigation will be required to dissect the functional contributions of the closed and open states of UPF1 to NMD.

An open question not addressed in our work is whether UPF1 dissociation from RNAs is inhibited at a particular stage of NMD, either before or after the commitment to decay. One possibility is that UPF1 phosphorylation and/or translation termination directly affect UPF1 dissociation rates by modulating ATP hydrolysis (5, 11). Alternatively, higher-order complex assembly initiated by SMG5, SMG6, and/or SMG7 binding to phosphorylated UPF1 may establish multivalent interactions capable of holding UPF1 in proximity to RNA despite the disruptive effect of ATP hydrolysis on UPF1-RNA interactions (14, 15). In addition, RNA-binding proteins such as staufen and the histone stem-loop binding protein may tether UPF1 to RNA, preventing its dissociation (50, 51). Similarly, the mammalian “long-loop” alternative isoform of UPF1 is less susceptible to ATPase-stimulated dissociation, allowing it to function on RNAs normally immune to decay (34).

## Acknowledgements

We thank members of the Hogg lab, Ian Morgan, Gabor Harami, and Sarah Fritz for critical discussion and reading of the manuscript. This work was supported by the Intramural Research Program, National Heart, Lung, and Blood Institute, National Institutes of Health, and utilized the computational resources of the NIH HPC Biowulf cluster (http://hpc.nih.gov). We would like to thank the NHLBI Biophysics core for training and use of the BLI and MST instruments and the NHLBI DNA Sequencing and Genomics core for providing NGS services.

## Author contributions

J.H.C., A.M.Y., K.C.N., and J.R.H. conceptualized the study. J.H.C. wrote the manuscript and all other authors edited it. A.M.Y. acquired and analyzed the majority of the *in vitro* experiments and cellular experiments. J.H.C. and A.L.G. acquired and analyzed initial *in vitro* experiments, and J.H.C. performed the *in silico* screen, computational modeling, RNA-seq experiments, and assisted in data analysis.

## References

1. Palumbo, M.C., Farina, L. and Paci, P. (2015) Kinetics effects and modeling of mRNA turnover. Wiley Interdiscip. Rev. RNA, 6, 327–336.

2. Kishor, A., Fritz, S.E. and Hogg, J.R. (2019) Nonsense-mediated mRNA decay: The challenge of telling right from wrong in a complex transcriptome. Wiley Interdiscip. Rev. RNA, 10, e1548.

3. Kim, Y.K. and Maquat, L.E. (2019) UPFront and center in RNA decay: UPF1 in nonsense-mediated mRNA decay and beyond. RNA, 25, 407–422.

4. Sun, L., Mailliot, J. and Schaffitzel, C. (2023) Nonsense-Mediated mRNA Decay Factor Functions in Human Health and Disease. Biomedicines, 11, 722.

5. Lee, S.R., Pratt, G.A., Martinez, F.J., Yeo, G.W. and Lykke-Andersen, J. (2015) Target Discrimination in Nonsense-Mediated mRNA Decay Requires Upf1 ATPase Activity. Mol. Cell, 59, 413–425.

6. Karousis, E.D. and Mühlemann, O. (2019) Nonsense-Mediated mRNA Decay Begins Where Translation Ends. Cold Spring Harb. Perspect. Biol., 11.

7. Hogg, J.R. and Goff, S.P. (2010) Upf1 senses 3’UTR length to potentiate mRNA decay. Cell, 143, 379–389.

8. He, F., Brown, A.H. and Jacobson, A. (1997) Upf1p, Nmd2p, and Upf3p are interacting components of the yeast nonsense-mediated mRNA decay pathway. Mol. Cell. Biol., 17, 1580–1594.

9. Serin, G., Gersappe, A., Black, J.D., Aronoff, R. and Maquat, L.E. (2001) Identification and characterization of human orthologues to Saccharomyces cerevisiae Upf2 protein and Upf3 protein (Caenorhabditis elegans SMG-4). Mol. Cell. Biol., 21, 209–223.

10. Clerici, M., Mourão, A., Gutsche, I., Gehring, N.H., Hentze, M.W., Kulozik, A., Kadlec, J., Sattler, M. and Cusack, S. (2009) Unusual bipartite mode of interaction between the nonsense-mediated decay factors, UPF1 and UPF2. EMBO J., 28, 2293–2306.

11. Czaplinski, K., Ruiz-Echevarria, M.J., Paushkin, S.V., Han, X., Weng, Y., Perlick, H.A., Dietz, H.C., Ter-Avanesyan, M.D. and Peltz, S.W. (1998) The surveillance complex interacts with the translation release factors to enhance termination and degrade aberrant mRNAs. Genes Dev., 12, 1665–1677.

12. Kashima, I., Yamashita, A., Izumi, N., Kataoka, N., Morishita, R., Hoshino, S., Ohno, M., Dreyfuss, G. and Ohno, S. (2006) Binding of a novel SMG-1-Upf1-eRF1-eRF3 complex (SURF) to the exon junction complex triggers Upf1 phosphorylation and nonsense-mediated mRNA decay. Genes Dev., 20, 355–367.

13. Ivanov, P.V., Gehring, N.H., Kunz, J.B., Hentze, M.W. and Kulozik, A.E. (2008) Interactions between UPF1, eRFs, PABP and the exon junction complex suggest an integrated model for mammalian NMD pathways. EMBO J., 27, 736–747.

14. Ohnishi, T., Yamashita, A., Kashima, I., Schell, T., Anders, K.R., Grimson, A., Hachiya, T., Hentze, M.W., Anderson, P. and Ohno, S. (2003) Phosphorylation of hUPF1 induces formation of mRNA surveillance complexes containing hSMG-5 and hSMG-7. Mol. Cell, 12, 1187–1200.

15. Okada-Katsuhata, Y., Yamashita, A., Kutsuzawa, K., Izumi, N., Hirahara, F. and Ohno, S. (2012) N- and C-terminal Upf1 phosphorylations create binding platforms for SMG-6 and SMG-5:SMG-7 during NMD. Nucleic Acids Res., 40, 1251–1266.

16. Loh, B., Jonas, S. and Izaurralde, E. (2013) The SMG5-SMG7 heterodimer directly recruits the CCR4-NOT deadenylase complex to mRNAs containing nonsense codons via interaction with POP2. Genes Dev., 27, 2125–2138.

17. Nicholson, P., Josi, C., Kurosawa, H., Yamashita, A. and Mühlemann, O. (2014) A novel phosphorylation-independent interaction between SMG6 and UPF1 is essential for human NMD. Nucleic Acids Res., 42, 9217–9235.

18. Chakrabarti, S., Bonneau, F., Schüssler, S., Eppinger, E. and Conti, E. (2014) Phospho-dependent and phospho-independent interactions of the helicase UPF1 with the NMD factors SMG5-SMG7 and SMG6. Nucleic Acids Res., 42, 9447–9460.

19. Czaplinski, K., Weng, Y., Hagan, K.W. and Peltz, S.W. (1995) Purification and characterization of the Upf1 protein: a factor involved in translation and mRNA degradation. RNA, 1, 610– 623.

20. Weng, Y., Czaplinski, K. and Peltz, S.W. (1996) Genetic and biochemical characterization of mutations in the ATPase and helicase regions of the Upf1 protein. Mol. Cell. Biol., 16, 5477–5490.

21. Fiorini, F., Bagchi, D., Le Hir, H. and Croquette, V. (2015) Human Upf1 is a highly processive RNA helicase and translocase with RNP remodelling activities. Nat. Commun., 6, 7581.

22. Serdar, L.D., Whiteside, D.L. and Baker, K.E. (2016) ATP hydrolysis by UPF1 is required for efficient translation termination at premature stop codons. Nat. Commun., 7, 14021.

23. Serdar, L.D., Whiteside, D.L., Nock, S.L., McGrath, D. and Baker, K.E. (2020) Inhibition of post-termination ribosome recycling at premature termination codons in UPF1 ATPase mutants. Elife, 9.

24. Chakrabarti, S., Jayachandran, U., Bonneau, F., Fiorini, F., Basquin, C., Domcke, S., Le Hir, H. and Conti, E. (2011) Molecular mechanisms for the RNA-dependent ATPase activity of Upf1 and its regulation by Upf2. Mol. Cell, 41, 693–703.

25. Kadlec, J., Guilligay, D., Ravelli, R.B. and Cusack, S. (2006) Crystal structure of the UPF2-interacting domain of nonsense-mediated mRNA decay factor UPF1. RNA, 12, 1817–1824.

26. Melero, R., Uchiyama, A., Castaño, R., Kataoka, N., Kurosawa, H., Ohno, S., Yamashita, A. and Llorca, O. (2014) Structures of SMG1-UPFs complexes: SMG1 contributes to regulate UPF2-dependent activation of UPF1 in NMD. Structure, 22, 1105–1119.

27. Deniaud, A., Karuppasamy, M., Bock, T., Masiulis, S., Huard, K., Garzoni, F., Kerschgens, K., Hentze, M.W., Kulozik, A.E., Beck, M., et al. (2015) A network of SMG-8, SMG-9 and SMG-1 C-terminal insertion domain regulates UPF1 substrate recruitment and phosphorylation. Nucleic Acids Res., 43, 7600–7611.

28. Langer, L.M., Gat, Y., Bonneau, F. and Conti, E. (2020) Structure of substrate-bound SMG1-8-9 kinase complex reveals molecular basis for phosphorylation specificity. Elife, 9.

29. Xue, G., Maciej, V.D., Machado de Amorim, A., Pak, M., Jayachandran, U. and Chakrabarti, S. (2022) Modulation of RNA binding properties of the RNA helicase UPF1 by its activator UPF2. RNA, 10.1261/rna.079188.122.

30. Weng, Y., Czaplinski, K. and Peltz, S.W. (1998) ATP is a cofactor of the Upf1 protein that modulates its translation termination and RNA binding activities. RNA, 4, 205–214.

31. Chapman, J.H., Craig, J.M., Wang, C.D., Gundlach, J.H., Neuman, K.C. and Hogg, J.R. (2022) UPF1 mutants with intact ATPase but deficient helicase activities promote efficient nonsense-mediated mRNA decay. Nucleic Acids Res., 10.1093/nar/gkac1026.

32. Fritz, S.E., Ranganathan, S., Wang, C.D. and Hogg, J.R. (2020) The RNA-binding protein PTBP1 promotes ATPase-dependent dissociation of the RNA helicase UPF1 to protect transcripts from nonsense-mediated mRNA decay. J. Biol. Chem., 295, 11613–11625.

33. Durand, S., Franks, T.M. and Lykke-Andersen, J. (2016) Hyperphosphorylation amplifies UPF1 activity to resolve stalls in nonsense-mediated mRNA decay. Nat. Commun., 7, 12434.

34. Fritz, S.E., Ranganathan, S., Wang, C.D. and Hogg, J.R. (2022) An alternative UPF1 isoform drives conditional remodeling of nonsense-mediated mRNA decay. EMBO J., 41, e108898.

35. Kurosaki, T., Li, W., Hoque, M., Popp, M.W.-L., Ermolenko, D.N., Tian, B. and Maquat, L.E. (2014) A post-translational regulatory switch on UPF1 controls targeted mRNA degradation. Genes Dev., 28, 1900–1916.

36. Pettersen, E.F., Goddard, T.D., Huang, C.C., Couch, G.S., Greenblatt, D.M., Meng, E.C. and Ferrin, T.E. (2004) UCSF Chimera--a visualization system for exploratory research and analysis. J. Comput. Chem., 25, 1605–1612.

37. Fritz, S.E., Haque, N. and Hogg, J.R. (2018) Highly efficient in vitro translation of authentic affinity purified messenger ribonucleoprotein complexes. RNA, 10.1261/rna.065730.118.

38. Baldwin, A., Morris, A.R. and Mukherjee, N. (2021) An Easy, Cost-Effective, and Scalable Method to Deplete Human Ribosomal RNA for RNA-seq. Curr Protoc, 1, e176.

39. Chamieh, H., Ballut, L., Bonneau, F. and Le Hir, H. (2008) NMD factors UPF2 and UPF3 bridge UPF1 to the exon junction complex and stimulate its RNA helicase activity. Nat. Struct. Mol. Biol., 15, 85–93.

40. Gowravaram, M., Bonneau, F., Kanaan, J., Maciej, V.D., Fiorini, F., Raj, S., Croquette, V., Le Hir, H. and Chakrabarti, S. (2018) A conserved structural element in the RNA helicase UPF1 regulates its catalytic activity in an isoform-specific manner. Nucleic Acids Res., 46, 2648– 2659.

41. Gowravaram, M., Schwarz, J., Khilji, S.K., Urlaub, H. and Chakrabarti, S. (2019) Insights into the assembly and architecture of a Staufen-mediated mRNA decay (SMD)-competent mRNP. Nat. Commun., 10, 5054.

42. Franks, T.M., Singh, G. and Lykke-Andersen, J. (2010) Upf1 ATPase-dependent mRNP disassembly is required for completion of nonsense-mediated mRNA decay. Cell, 143, 938–950.

43. Cho, N.H., Cheveralls, K.C., Brunner, A.-D., Kim, K., Michaelis, A.C., Raghavan, P., Kobayashi, H., Savy, L., Li, J.Y., Canaj, H., et al. (2022) OpenCell: Endogenous tagging for the cartography of human cellular organization. Science, 375, eabi6983.

44. Wang, M., Weiss, M., Simonovic, M., Haertinger, G., Schrimpf, S.P., Hengartner, M.O. and von Mering, C. (2012) PaxDb, a database of protein abundance averages across all three domains of life. Mol. Cell. Proteomics, 11, 492–500.

45. Cheng, Z., Muhlrad, D., Lim, M.K., Parker, R. and Song, H. (2007) Structural and functional insights into the human Upf1 helicase core. EMBO J., 26, 253–264.

46. Bhattacharya, A., Czaplinski, K., Trifillis, P., He, F., Jacobson, A. and Peltz, S.W. (2000) Characterization of the biochemical properties of the human Upf1 gene product that is involved in nonsense-mediated mRNA decay. RNA, 6, 1226–1235.

47. Leeds, P., Wood, J.M., Lee, B.S. and Culbertson, M.R. (1992) Gene products that promote mRNA turnover in Saccharomyces cerevisiae. Mol. Cell. Biol., 12, 2165–2177.

48. Sun, X., Perlick, H.A., Dietz, H.C. and Maquat, L.E. (1998) A mutated human homologue to yeast Upf1 protein has a dominant-negative effect on the decay of nonsense-containing mRNAs in mammalian cells. Proc. Natl. Acad. Sci. U. S. A., 95, 10009–10014.

49. Kanaan, J., Raj, S., Decourty, L., Saveanu, C., Croquette, V. and Le Hir, H. (2018) UPF1-like helicase grip on nucleic acids dictates processivity. Nat. Commun., 9, 3752.

50. Kim, Y.K., Furic, L., DesGroseillers, L. and Maquat, L.E. (2005) Mammalian Staufen1 recruits Upf1 to specific mRNA 3′ UTRs so as to elicit mRNA decay. Cell, 120, 195–208.

51. Kaygun, H. and Marzluff, W.F. (2005) Regulated degradation of replication-dependent histone mRNAs requires both ATR and Upf1. Nat. Struct. Mol. Biol., 12, 794–800.

52. Waskom, M. (2021) seaborn: statistical data visualization. J. Open Source Softw., 6, 3021.

53. Virtanen, P., Gommers, R., Oliphant, T.E., Haberland, M., Reddy, T., Cournapeau, D., Burovski, E., Peterson, P., Weckesser, W., Bright, J., et al. (2020) SciPy 1.0: fundamental algorithms for scientific computing in Python. Nat. Methods, 17, 261–272.

54. Martin, M. (2011) Cutadapt removes adapter sequences from high-throughput sequencing reads. EMBnet.journal, 17, 10–12.

55. Dobin, A., Davis, C.A., Schlesinger, F., Drenkow, J., Zaleski, C., Jha, S., Batut, P., Chaisson, M. and Gingeras, T.R. (2013) STAR: ultrafast universal RNA-seq aligner. Bioinformatics, 29, 15–21.

56. Li, B. and Dewey, C.N. (2011) RSEM: accurate transcript quantification from RNA-Seq data with or without a reference genome. BMC Bioinformatics, 12, 323.

57. Love, M.I., Huber, W. and Anders, S. (2014) Moderated estimation of fold change and dispersion for RNA-seq data with DESeq2. Genome Biol., 15, 550.

58. Ge, S.X., Son, E.W. and Yao, R. (2018) iDEP: an integrated web application for differential expression and pathway analysis of RNA-Seq data. BMC Bioinformatics, 19, 534.

59. Shen, S., Park, J.W., Lu, Z.-X., Lin, L., Henry, M.D., Wu, Y.N., Zhou, Q. and Xing, Y. (2014) rMATS: Robust and flexible detection of differential alternative splicing from replicate RNA-Seq data. Proceedings of the National Academy of Sciences, 111, E5593–E5601.

