## Supplemental Material for "UPF1 ATPase autoinhibition and activation modulate RNA binding kinetics and NMD efficiency"

**Supplementary Material**

**Supplementary Tables**

**Supplementary Table S1.** Oligonucleotides used for *in vitro* assays and primers used for generating UPF1 and UPF2 constructs

**Supplementary Table S2.** RT-qPCR Primers

**Supplementary Table S3.** Oligonucleotides used for rRNA depletion

**Supplementary Table S4.** Kinetic parameters used for *in silico* modeling of UPF2-UPF1-RNA interactions

**Supplementary Table S5.** Results of *in silico* Rosetta-Vienna screen

**Supplementary Table S6.** Comparison of *in silico* Rosetta-Vienna screens between UPF1-CH-HD and UPF1-HD

**Supplementary Table S7.** RT-qPCR Statistics

**Supplementary Table S8.** RNA-seq data analysis

**Supplementary Materials & Methods**

***Fitting and calculations for Biolayer interferometry (BLI) and fluorescence anisotropy dissociation (FAD) assays***

To obtain ATP-dependent off-rates in both BLI and FAD experiments, curves were fit to double exponential decay functions ($e^{-kt}$), and $k_1$ was reported as the ATP-dependent off-rate as it was always weighted higher than $k_2$. For on-rates in BLI experiments, each curve was fit to a double exponential growth curve ($1 - e^{-kt}$), and $k_1$ was used in subsequent downstream analyses as it was always weighted higher than $k_2$. $k_1$ values were plotted for each concentration, and the slope of this curve was extracted to calculate $k_{on}$ in units of $nM^{-1}sec^{-1}$.

***Processivity calculations/simulations, unwinding data fitting, unwinding activity calculations***

All fitting, calculations, and simulations were performed as previously described (31). To calculate unwinding activity from multiple turnover fluorescence-based unwinding assays, all fluorescence values were first normalized so that the starting fluorescence was 1. Areas under +ATP curves (normalized to the corresponding -ATP curve to account for any drift) were then estimated using Simpson's rule, implemented using the scipy.integrate.simpson python package (53). Areas were subtracted from the final time point of the data (generally 1200), and this number was divided by the final time point of the data to generate the "unwinding activity" parameter, which accounts for both unwinding rate and processivity and is a number between zero and one. For example, no unwinding would give an area of 1200, so "unwinding activity" would equal zero. Conversely, a rapid drop in fluorescence from one to zero within the first time point would give an area of zero, so "unwinding activity" would equal one.

***In silico modeling of UPF1-UPF2-RNA interactions***

All modeling was performed using agent-based computational modeling with custom python scripts. Each UPF1, UPF2, and RNA object was given experimentally determined parameters based on FAD and BLI kinetic assays (see **Supplementary Table S4**). UPF1 on-rates were multiplied by 1000 to account for the approximate 1000 nM UPF1 concentration in cells, and UPF2 on-rates were multiplied by 100 to account for the approximate 100 nM concentration in cells. Bimolecular interactions did not need to be considered given the relative cellular concentrations. To obtain probabilities of binding and dissociation from these rates, the exponential distribution needs to first be normalized. For example, with a rate of 0.08 $sec^{-1}$:

$$\frac{1}{\int_0^\infty e^{-0.08t} dt} = 0.08$$

Next, the probability is calculated between 0 and the time step (1 second in all simulations performed) to get the probability of either binding or dissociating each time step:

$$0.08 \int_0^1 e^{-0.08t} dt = 0.0769$$

Simulations were initiated by pre-binding every UPF1 object to a different RNA object, such that all 1000 RNA objects each had one unique UPF1 object bound. Each time step (representing one second), each UPF2 object was given a chance to bind or unbind from a UPF1 object based on the kinetic parameters extracted from BLI experiments. Also at each time step, each UPF1 object was given a chance to bind or unbind from an RNA object based on the kinetic parameters extracted from BLI and FAD experiments. If a UPF1 object was not UPF2-bound, the parameters were taken from UPF1-CH-HD WT, but if a UPF1 object was UPF2-bound, the parameters were taken from the average of UPF1-CH-HD F192E and I757R. At the end of each time step, the fraction of UPF1 objects bound to UPF2, bound to RNA, bound to both, or unbound was calculated. Also at each time step, the number of RNA binding and dissociation steps were counted and added to a cumulative total.

### *In silico modeling of UPF1 unwinding RNA under single turnover fluorescence-based unwinding assay conditions*

All simulations were performed using agent-based computational modeling with custom python scripts. All simulated UPF1 molecules were given the intrinsic properties of processivity, unwinding rate in the open state, unwinding rate in the closed state, probability of transitioning to the open state, and transitioning to the closed state. All simulated RNA molecules had a length of 42 nt (corresponding to a 10 bp hpDNA used in experiential unwinding assays) and a starting fluorescence of 1. Simulations started with binding of all UPF1 molecules, such that every RNA contained one unique UPF1 bound at the 5′ end. At each bp step, UPF1 could transition to the open or closed state, with the outcome determined by comparing the intrinsic probability to a random flat probability distribution between 0 and 1. Furthermore, for each bp step, the time to take the step was drawn from an exponential distribution corresponding to the unwinding rate. The closed and open states were modeled to have different unwinding rates. If UPF1 stepped 42 times without falling off (dependent on processivity), the fluorescence of the RNA would change from 1 to 0 and the time to take 42 steps would be recorded. If a dissociation event occurred, the UPF1 molecule was no longer able to bind RNA, and the RNA it dissociated from would subsequently maintain its fluorescence of 1 throughout the simulation.

### *Gene-level analysis of RNA-seq data*

Workflow management programs CCBR Pipeliner (Figure 5D and Supplementary Figure S11A; https://github.com/CCBR/Pipeliner) and RENEE (Figure 6A; https://github.com/CCBR/RENEE) were used to analyze high-throughput sequencing data on the NIH Biowulf high-performance computing system. FASTQ files were trimmed to remove TruSeq adapter sequences and quality filtered using Cutadapt (54), with parameters --pair-filter=any --nextseq-trim=2 --trim-n -n 5 -O 5 -q 10,10 -m 35:35. Trimmed reads were subjected to two-pass STAR (55) alignment. First-pass STAR parameters were --outFilterIntronMotifs RemoveNoncanonicalUnannotated --outSAMstrandField None --outFilterType BySJout --outFilterMultimapNmax 20 --alignSJoverhangMin 8 --alignSJDBoverhangMin 1 --outFilterMismatchNmax 999 --outFilterMismatchNoverLmax 0.3 --alignIntronMin 20 --alignIntronMax 1000000 --alignMatesGapMax 1000000 --clip3pAdapterSeq - --outSAMtype BAM Unsorted --alignEndsProtrude 10 ConcordantPair --peOverlapNbasesMin 10. Second-pass STAR parameters were --outFilterIntronMotifs RemoveNoncanonicalUnannotated --outSAMstrandField None --outFilterType BySJout --outFilterMultimapNmax 20 --alignSJoverhangMin 8 --alignSJDBoverhangMin 1 --outFilterMismatchNmax 999 --outFilterMismatchNoverLmax 0.3 --alignIntronMin 20 --alignIntronMax 1000000 --alignMatesGapMax 1000000 --clip3pAdapterSeq - --outSAMunmapped Within --outWigType None --outWigStrand Stranded --limitSjdbInsertNsj 4000000 --quantMode TranscriptomeSAM GeneCounts --outSAMtype BAM SortedByCoordinate --alignEndsProtrude 10 ConcordantPair --peOverlapNbasesMin 10. Gene-level transcript abundance was determined using RSEM quantification against Gencode transcript annotations (56), with parameters rsem-calculate-expression --no-bam-output --calc-ci --seed 12345 --bam --paired-end --forward-prob=$fp --estimate-rspd. Gene counts were passed through a CPM filter of 0.5 and subjected to differential expression analysis using DESeq2 (57), implemented in the CCBR Pipeliner workflow (Figure 5D and Supplementary Figure S11A) or on the iDEP web server (Figure 6A) (58). For both datasets, the resulting datatables were filtered so that rows (genes) containing an empty value in any cell were removed, then filtered for genes upregulated in the siUPF1 CLIP condition by the indicated $\log_2$ fold changes of the si UPF1 CLIP condition compared to the siNT CLIP condition (with FDR < 0.05). Where indicated, the datatable was further filtered for genes that were rescued in the siUPF1 WT condition determined by the following formula:

$$\text{rescue efficiency} = \frac{(siUPF1\ CLIP\ vs\ siNT\ CLIP\ fold\ change) - (siUPF1\ WT\ vs\ siNT\ CLIP\ fold\ change)}{(siUPF1\ CLIP\ vs\ siNT\ CLIP\ fold\ change) - (siNT\ CLIP\ vs\ siNT\ CLIP\ fold\ change)}$$

Custom python scripts were used to create dendrograms.

***Junction-level analysis of RNA-seq data using rMATS to determine exon skipping and inclusion events***

The BAM files resulting from the two-pass STAR alignment to the human genome were used as inputs into rMATS 4.1.0 software (59). Gencode gene annotations and parameters -t paired --readLength 150 --variable-read-length --libType fr-firststrand were used for rMATS analysis. rMATS analyses of cassette exon splicing were filtered to include only events represented by an average of 20 junction-spanning reads in CLIP control siNT and siUPF1 samples. For genes represented by multiple events, only the event with the largest average number of junction-spanning reads was retained. Events significantly regulated upon UPF1 depletion from CLIP control lines (i.e. siUPF1 CLIP condition) were defined as having FDR < 0.05 and |IncLevelDifference| > 0.1. Rescued events were defined as those in which > 75% of the IncLevelDifference caused by siUPF1 treatment was reversed by expression of WT CLIP-UPF1, using the following formula:

rescue efficiency = $\frac{(siUPF1\ CLIP\ vs\ siNT\ CLIP\ IncLevelDifference) - (siUPF1\ WT\ vs\ siNT\ CLIP\ IncLevelDifference)}{(siUPF1\ CLIP\ vs\ siNT\ CLIP\ IncLevelDifference)}$

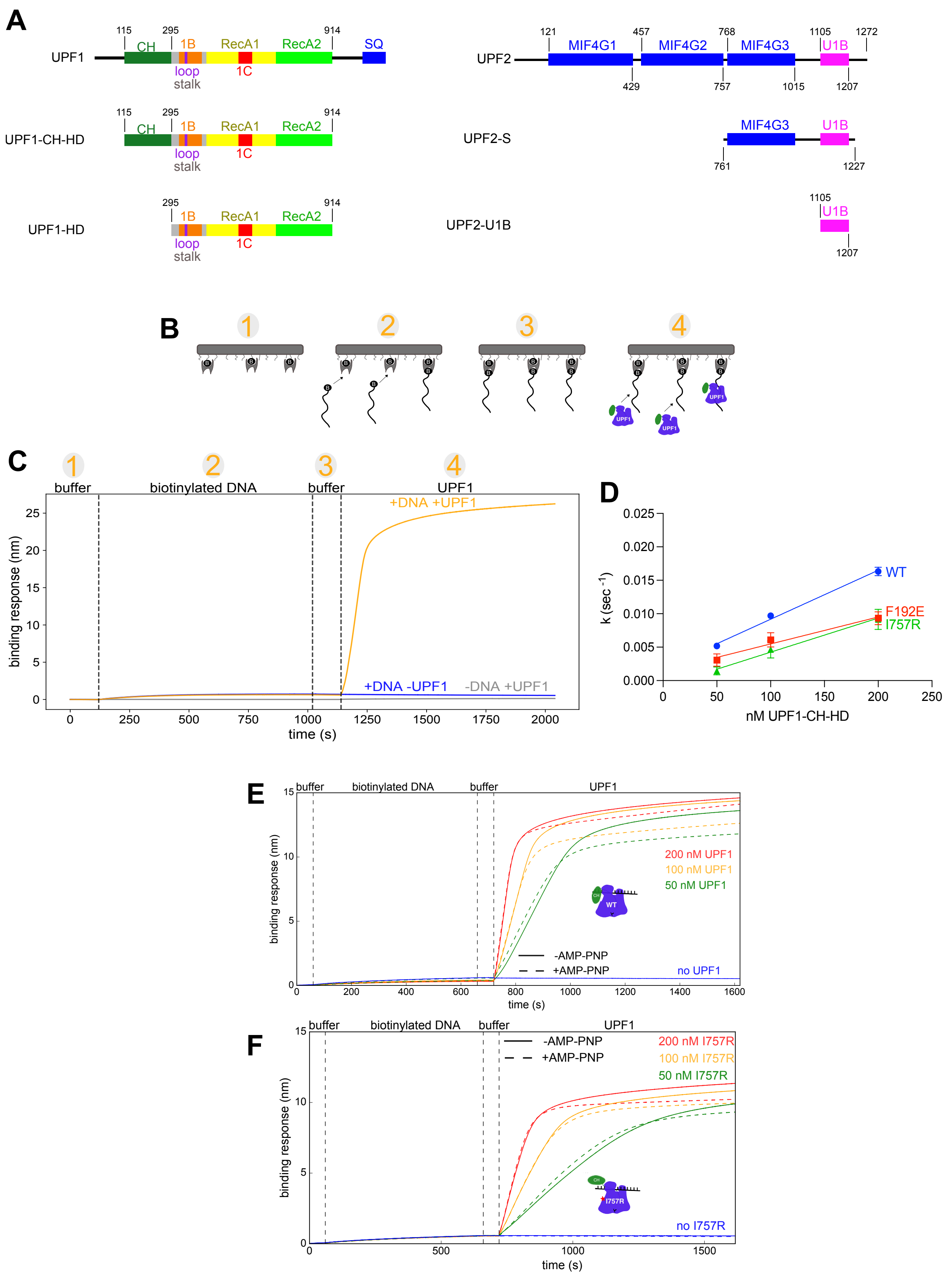


**Supplementary Figure S1: Related to Figure 1.**

***Supplementary Figure S1. UPF1-CH-HD exhibits specific binding to nucleic acid substrates in BLI experiments. Related to Figure 1.*** (**A**) Schematics of UPF1 (left) and UPF2 (right) constructs used in this study. CH is the autoinhibitory cysteine/histidine-rich domain. The 1B, 1C, and RecA domains comprise the UPF1 helicase domain. SQ is the domain containing several SQ motifs for phosphorylation. U1B is the UPF1 binding domain of UPF2. UPF1-CH-HD and UPF2-S were the default constructs for *in vitro* experiments unless otherwise specified. (**B**) Schematic of BLI as in Figure 1C. (**C**) Control BLI run of UPF1-CH-HD WT showing binding to the sensor surface in a DNA-dependent manner. There was no detectable non-specific surface binding (gray curve). (**D**) Extraction of k values for each concentration of UPF1-CH-HD from 4 independent experiments, in which the slope of the curves was calculated to extract the on-rates depicted in Figure 1F.

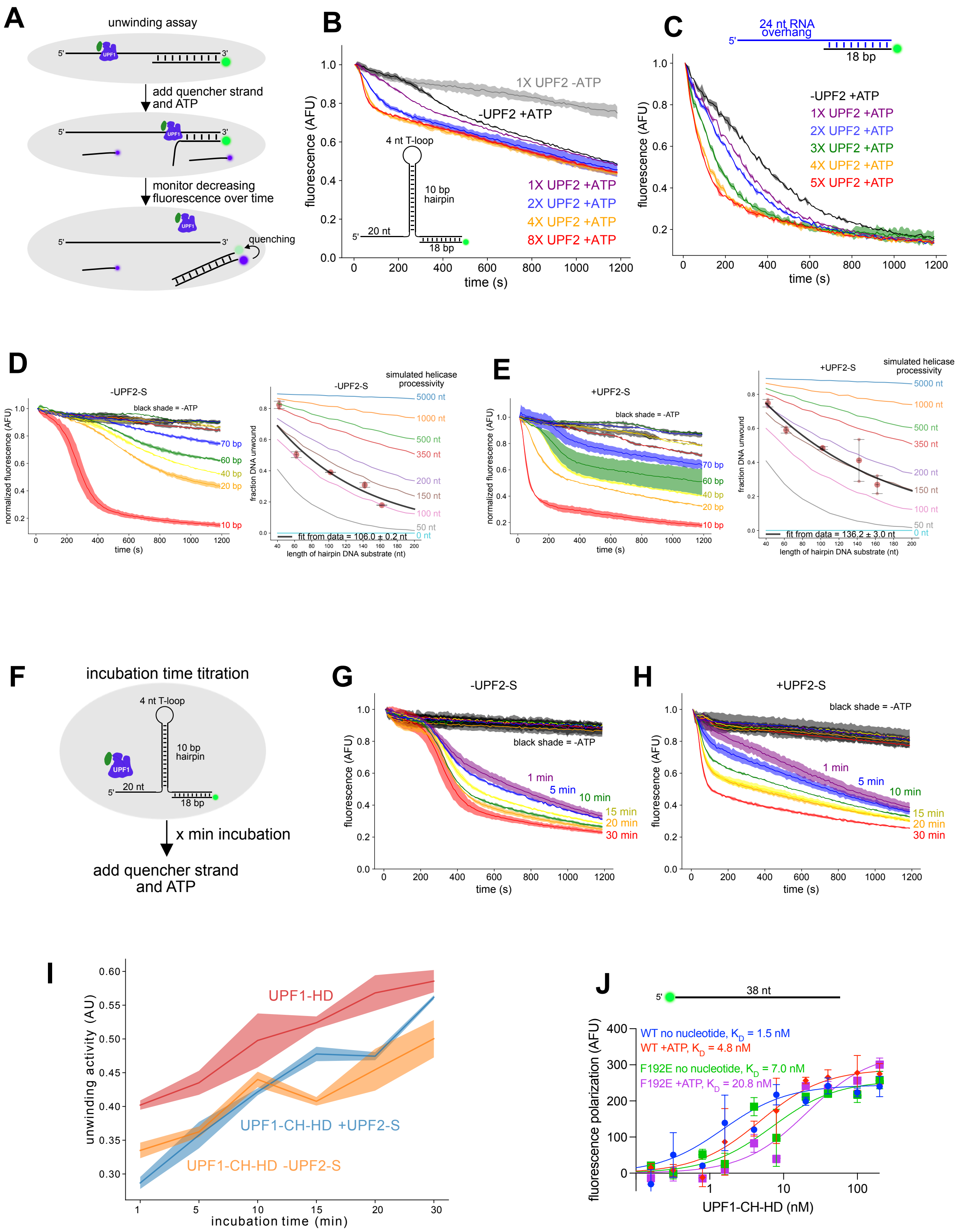


**Supplementary Figure S2: Related to Figure 1.**

***Supplementary Figure S2. UPF1-CH-HD activity but not processivity is enhanced by UPF2-S, and open UPF1 constructs are more sensitive to incubation time with the unwinding assay substrate. Related to Figure 1.*** (**A**) Multiple turnover unwinding assay schematic. UPF2-S was used in 2-fold molar excess (165 nM) to UPF1-CH-HD (82.5 nM) unless otherwise indicated. (**B**) Representative curves of UPF1-CH-HD unwinding hpDNA at varying concentrations of UPF2-S using the fluorescence-based unwinding assay. Shaded areas represent standard deviation. (**C**) As in (**B**) but with a substrate containing a 5′ RNA overhang. (**D**) Overlay of curves of UPF1-CH-HD multiple turnover unwinding with substrates as in (**B**) but with the indicated lengths of hairpin stem (left). Fraction of substrate unwound as a function of the hairpin stem length from 2 independent experiments (right). Black lines indicate processivity model fits and colored lines represent simulations of helicases with differing processivities (see **Supplementary Materials & Methods**). (**E**) As in (**D**) but with UPF2-S in the reaction from 2 independent experiments. (**F**) Schematic of multiple turnover unwinding assay with differing incubation times of the UPF1 construct with the 10 bp hpDNA substrate. (**G-H**) Representative curves of multiple turnover unwinding of UPF1-CH-HD in the absence (**G**) or presence of UPF2-S (**H**). Each curve indicates a different incubation time of UPF1 with the substrate. Complexes were formed by incubating UPF1 and UPF2 constructs at room temperature for > 10 min. (**I**) Calculation of unwinding activity (see **Supplementary Materials & Methods**) of each UPF1 construct at different incubation times from 3 independent experiments. Shaded regions represent standard deviation. (**J**) Equilibrium fluorescence anisotropy binding assay curves from 4 independent experiments in which UPF1-CH-HD constructs were incubated with 0.7 nM of a 38 nt ssDNA substrate for 2 hr in the presence and absence of ATP. Binding affinities ($K_D$) were calculated using the one-site specific binding nonlinear least squares fit in GraphPad Prism 9. Error bars represent standard deviation.

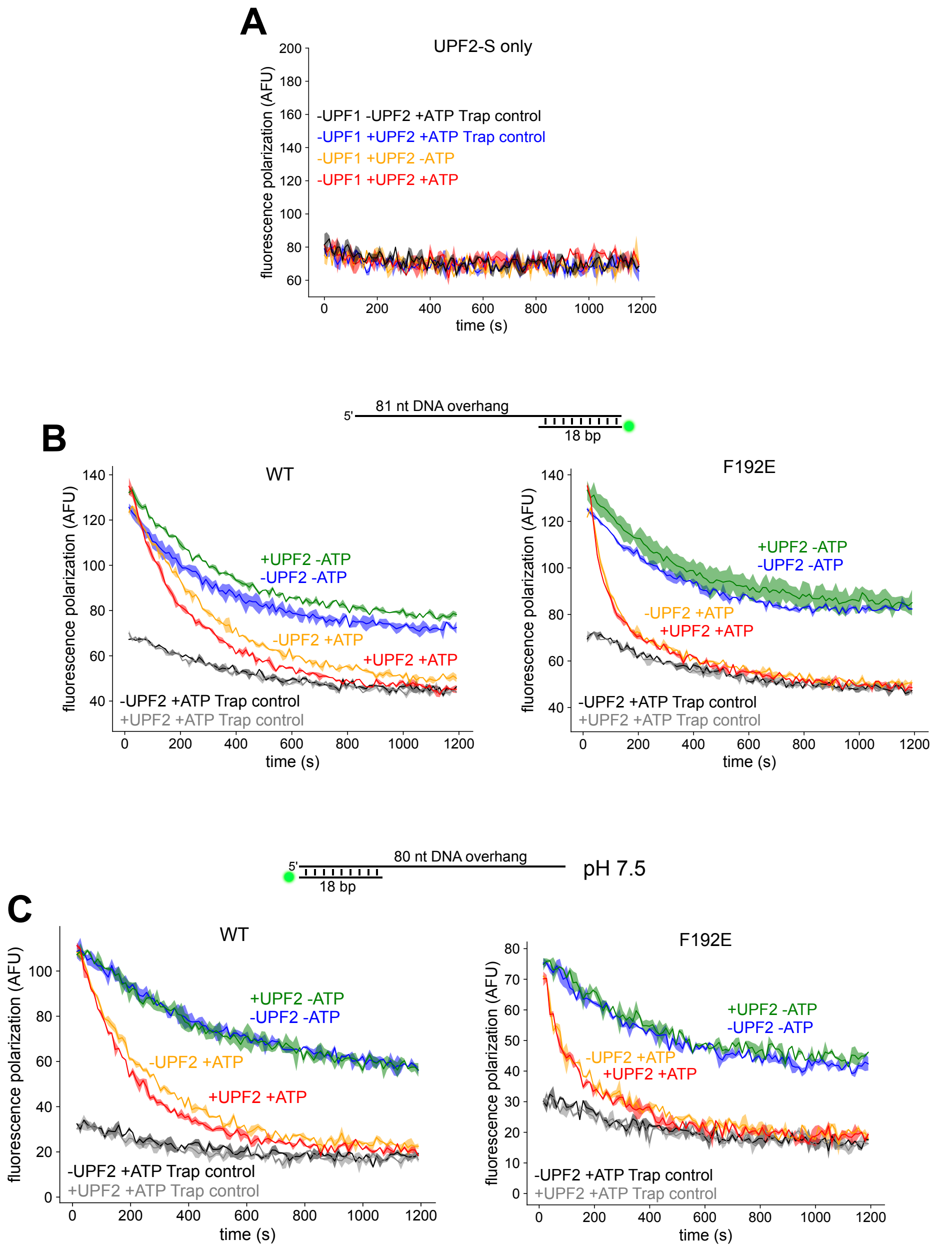


**Supplementary Figure S3: Related to Figure 2.**

***Supplementary Figure S3. The open state of UPF1 exhibits faster ATP-stimulated dissociation from nucleic acids. Related to Figure 2.*** (**A**) Representative curves of FAD with only UPF2-S to ensure no detectable binding of the fluorescent substrate from 2 independent experiments. (**B**) Representative curves of FAD of WT UPF1-CH-HD (left) or F192E (right) with an 81 nt 5′ DNA overhang substrate. 1000-fold molar excess of trap strand was added before UPF1-CH-HD addition to the substrate in trap controls (black and gray curves). (**C**) As in (**B**) but with an 80 nt 3′ DNA overhang substrate and under pH 7.5 instead of pH 6.0 conditions (see **Materials & Methods**).

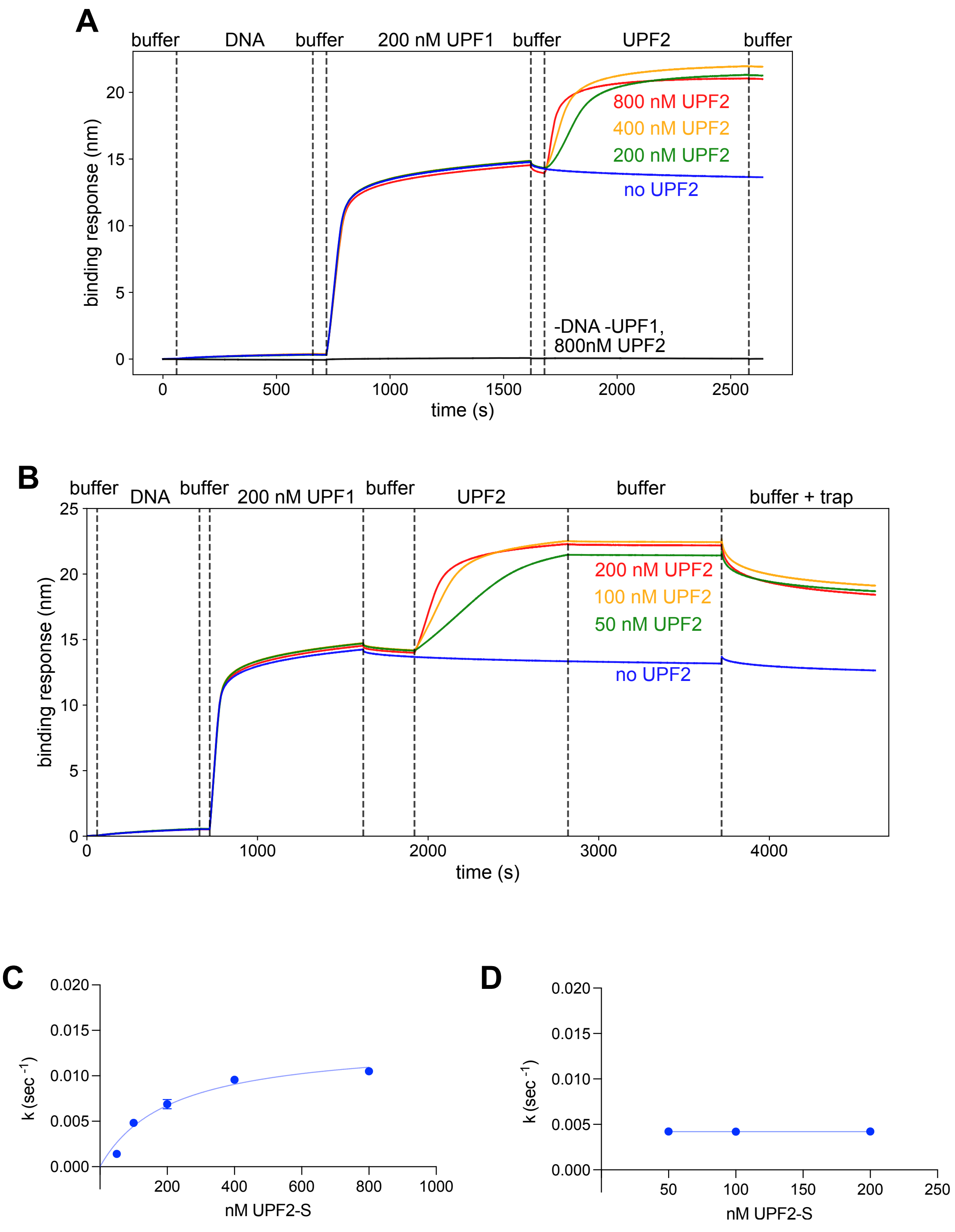


**Supplementary Figure S4: Related to Figure 3.**

***Supplementary Figure S4. BLI experiments of UPF2-S binding and dissociation from UPF1-CH-HD. Related to Figure 3.*** (**A**) Control BLI traces showing lack of UPF2-S non-specific binding to the surface (black curve) and specific binding to UPF1-CH-HD in a concentration-dependent manner. (**B**) As in (**A**) but with lower concentrations of UPF2-S and including dissociation steps. UPF2-S did not dissociate when sensors were dipped into buffer due to high affinity to UPF1-CH-HD, but modest dissociation occurred with 50 µM of 51 nt trap strand due to modest UPF2-S affinity to nucleic acids. (**C-D**) Extraction of UPF2-S on-rate to UPF1-CH-HD (**C**) and off-rate from UPF1-CH-HD (**D**) from 2 independent experiments.

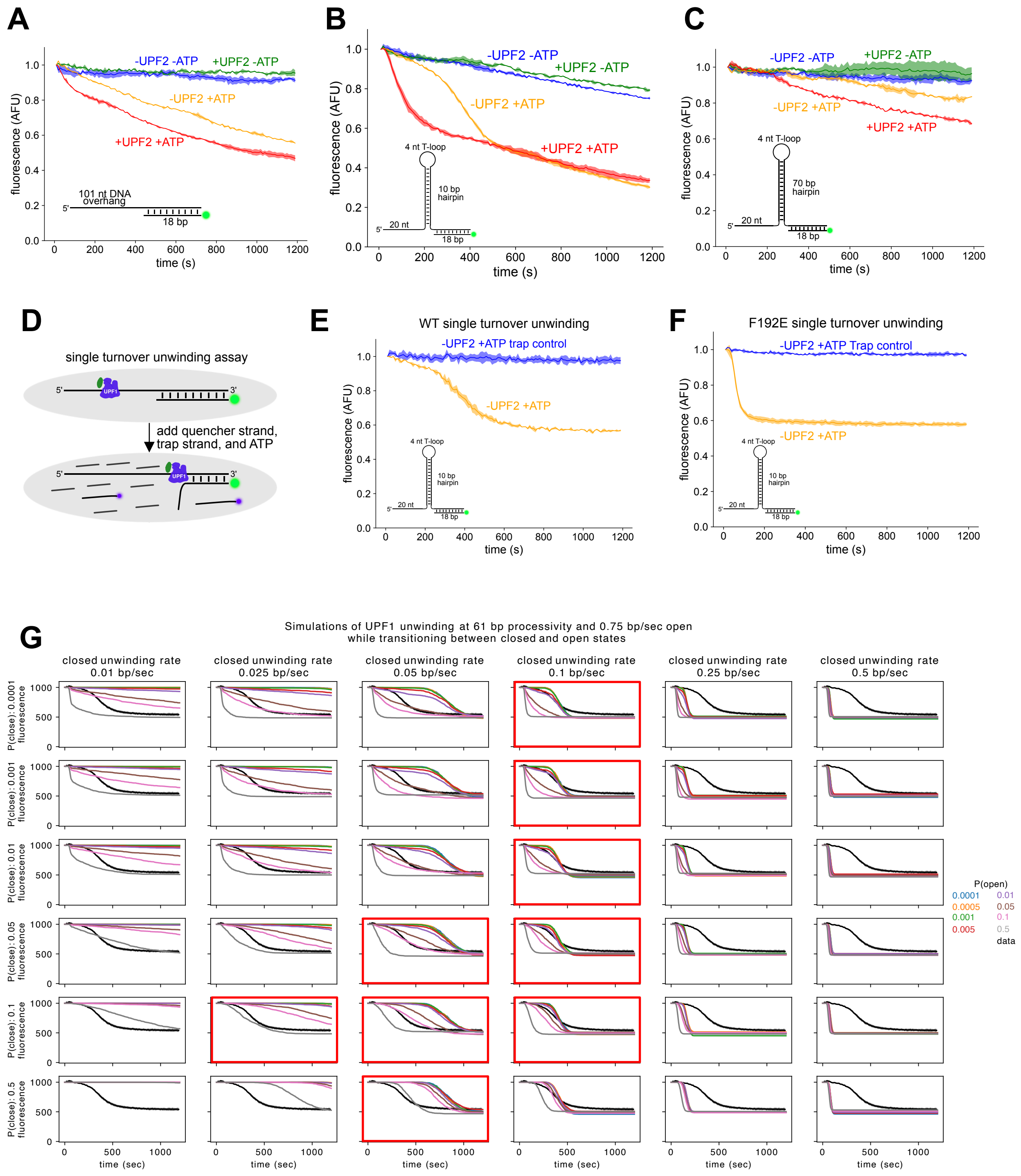


**Supplementary Figure S5: Related to Figure 3.**

***Supplementary Figure S5. In silico modeling of UPF1 unwinding at various parameters. Related to Figure 3.*** (**A-C**) Representative curves of UPF1-CH-HD unwinding a 101 nt 5′ DNA overhang substrate (**A**), short 10 bp hpDNA (**B**), or long 70 bp hpDNA (**C**) +/- UPF2-S. (**D**) Single turnover unwinding assay schematic. (**E-F**) Single turnover unwinding assay with UPF1-CH-HD WT (**E**) or F192E (**F**). 1000X of an unlabeled trap strand was added immediately before ATP addition (orange curves) or as a control before UPF1-CH-HD addition to the substrate (blue curves). (**G**) *In silico* modeling of 1000 UPF1s unwinding 1000 RNAs in a single turnover unwinding assay depicted in (**D**) to predict how different parameters of UPF1 unwinding would produce a biphasic curve shape. P(open) and P(close) are the probabilities of UPF1 opening or closing each bp step. Panels boxed in red indicate that at least one curve matches the 'data' curve. See **Supplementary Materials & Methods** for details.

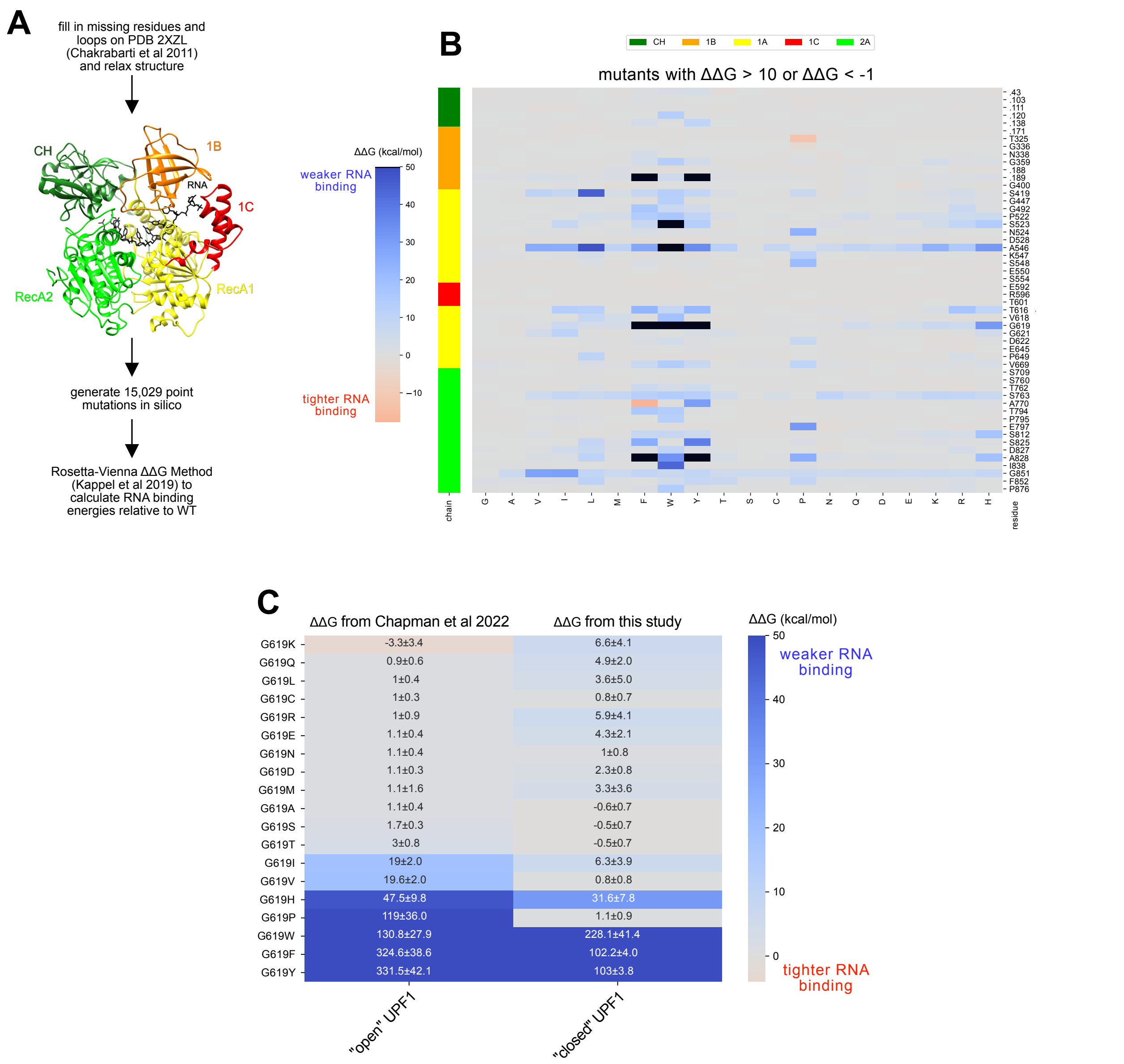


**Supplementary Figure S6: Related to Figure 4.**

***Supplementary Figure S6. Rosetta-Vienna ΔΔG Method identifies G619 as an important residue in the interaction between UPF1 and RNA in both the closed and open states. Related to Figure 4.*** (**A**) Computational workflow to identify mutations that affect UPF1-CH-HD RNA binding (see **Materials & Methods**) using PDB 2XZL (24). (**B**) Residues with at least 1 mutation of ΔΔG > 10 or ΔΔG < -1 (change in free binding energy compared to WT, units in kcal/mol) predicted by the *in silico* screen in (**A**). Predicted ΔΔG for each mutation was calculated by comparing the 20 lowest energy relaxed WT structures, and the average of the 20 comparisons is shown. Black boxes represent ΔΔG > 50. (**C**) Predicted ΔΔG values for each G619 mutation (right column) compared to the previous identical screen using the UPF1-HD structure (PDB 2XZO, left column) (31).

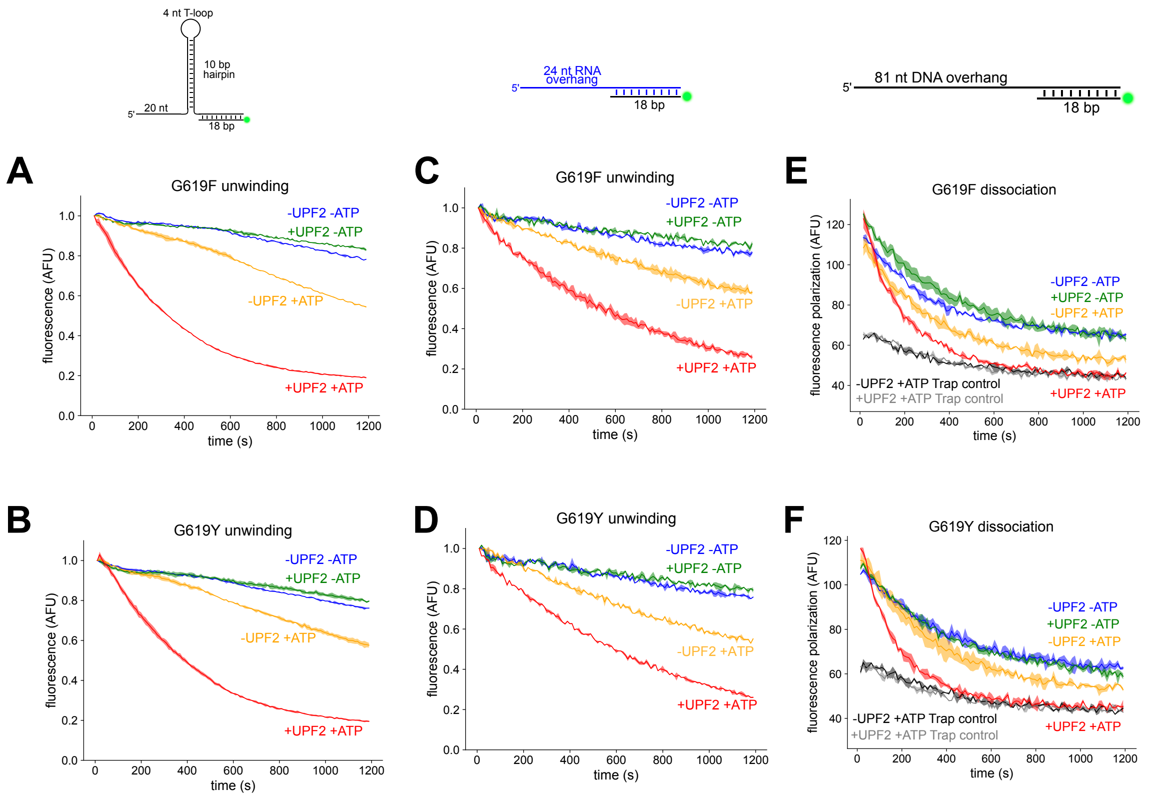


**Supplementary Figure S7: Related to Figure 4.**

***Supplementary Figure S7. Mutating UPF1-CH-HD G619 to an aromatic amino acid increases autoinhibition. Related to Figure 4.*** (**A-B**) Representative curves from multiple turnover unwinding with a short hpDNA substrate of UPF1-CH-HD G619F (**B**) or G619Y (**C**). Quantification of unwinding activity is shown in Figure 4D. (**C-D**) As in (**A-B**) but with a 5′ RNA overhang substrate and with 4-fold molar excess of UPF2-S (330 nM). (**G**) Quantification of unwinding activity is shown in Figure 4G. (**E-F**) Representative curves of FAD with a 5′ DNA overhang substrate of UPF1-CH-HD G619F (**E**) or G619Y (**F**). 1000-fold molar excess of trap strand was added before UPF1-CH-HD addition to the substrate in trap controls (black and gray curves). Half-lives calculated from FAD assays are shown in Figure 4J.

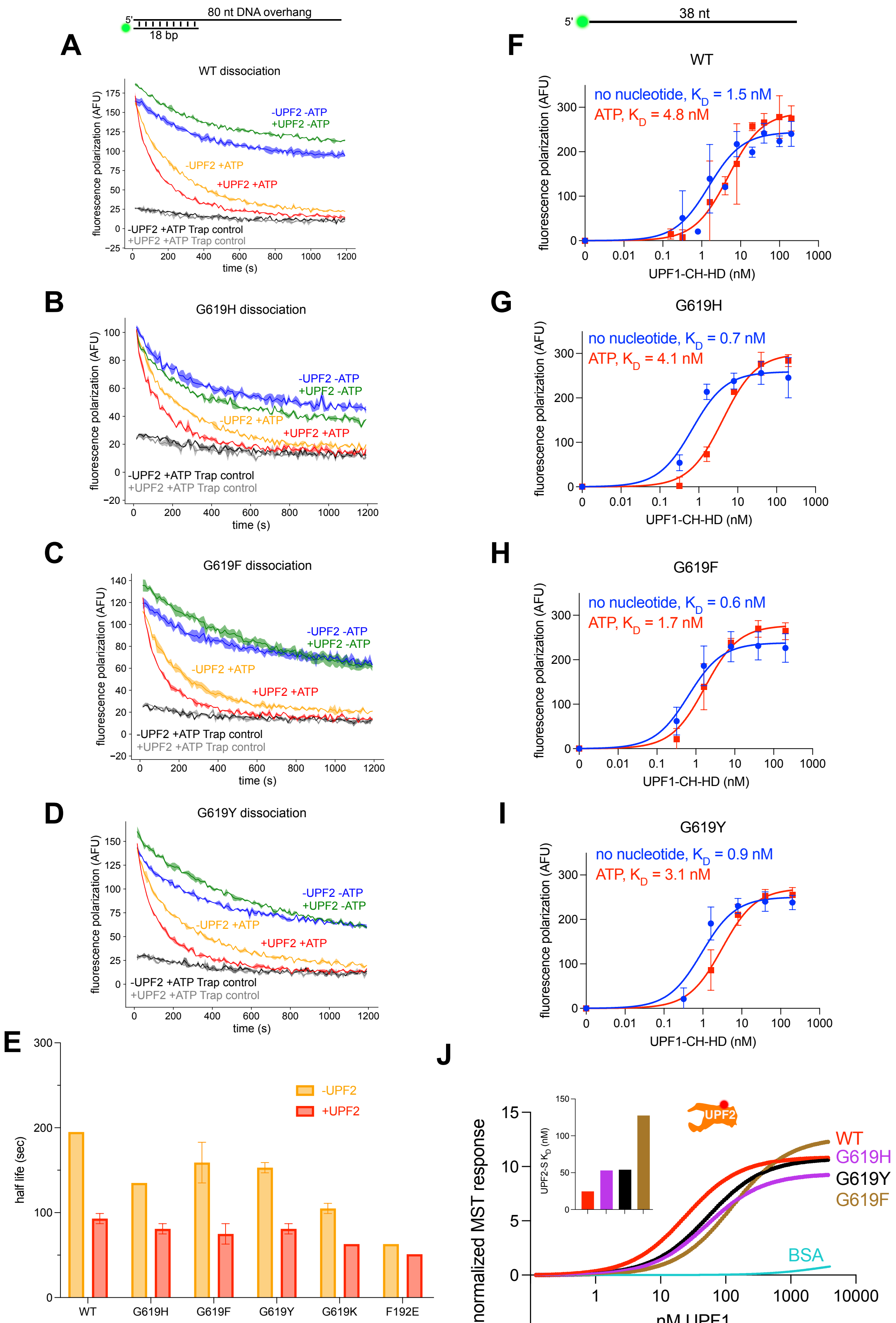


**Supplementary Figure S8: Related to Figure 4.**

***Supplementary Figure S8. More autoinhibited G619H, G619F, and G619Y UPF1-CH-HD mutants do not have nucleic acid or UPF2-S binding defects, or dissociation defects on 3′ overhang substrates. Related to Figure 4.*** (**A-D**) Representative curves of FAD with an 80 nt 3′ DNA overhang substrate of WT (**A**), G619H (**B**), G619F (**C**), or G619Y (**D**). (**E**) Calculation of half-lives from FAD assays as in Figure 4J but with an 80 nt 3′ DNA overhang substrate, from 2 independent experiments. (**F-I**) Equilibrium fluorescence anisotropy binding assay curves from 4 independent experiments in which the indicated UPF1-CH-HD constructs were incubated with a 38 nt ssDNA substrate for 2 hr +/- ATP. Binding affinities ($K_D$) were calculated using the one site specific binding nonlinear least squares fit in GraphPad Prism 9. Error bars represent standard deviation. (**J**) MST measurements of UPF1 binding to UPF2-S. UPF1 was titrated with 3.125 nM UPF2-S labeled with Alexa Fluor 647. Inset shows the UPF1 $K_D$ values for binding to UPF2-S after fitting curves. The BSA control did not show a sufficient MST response to calculate a $K_D$.

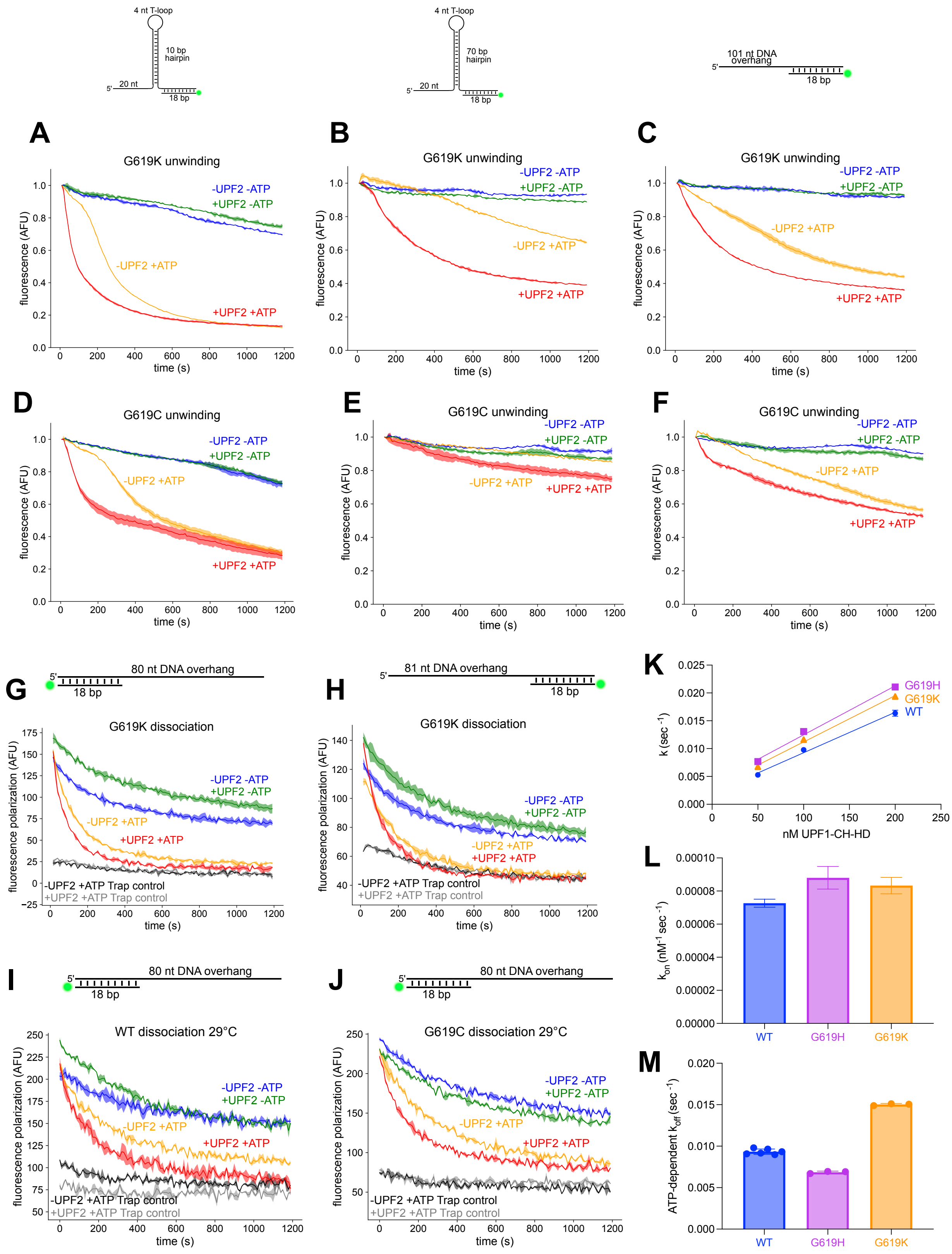


**Supplementary Figure S9: Related to Figure 4.**

***Supplementary Figure S9. Mutating residue G619 does not influence nucleic acid on-rate, and mutating G619 to a non-aromatic amino acid does not affect unwinding or dissociation activities. Related to Figure 4.*** (**A-F**) Representative curves of multiple turnover unwinding of UPF1-CH-HD G619K (**A-C**) or G619C (**D-F**) with 10 bp hpDNA (left graphs), 70 bp hpDNA (middle graphs), or 101 nt 5′ DNA overhang substrates (right graphs). (**G-H**) Representative curves of FAD with G619K on 3′ overhang DNA substrates (**G**) or 5′ overhang DNA substrates (**H**). (**I-J**) Representative curves of FAD at 29°C with WT (**I**) or G619C (**J**) on 3′ overhang DNA substrates. Unwinding activity calculations are shown in Figure 4D and 4G. Half life calculations are shown in Figure 4J. (**K**) Extraction of k values for each concentration of UPF1-CH-HD from 2 independent experiments. (**L**) Extraction of on-rates from the slope of the curves in (**K**). (**M**) ATP-dependent off-rates from 2 independent BLI experiments.

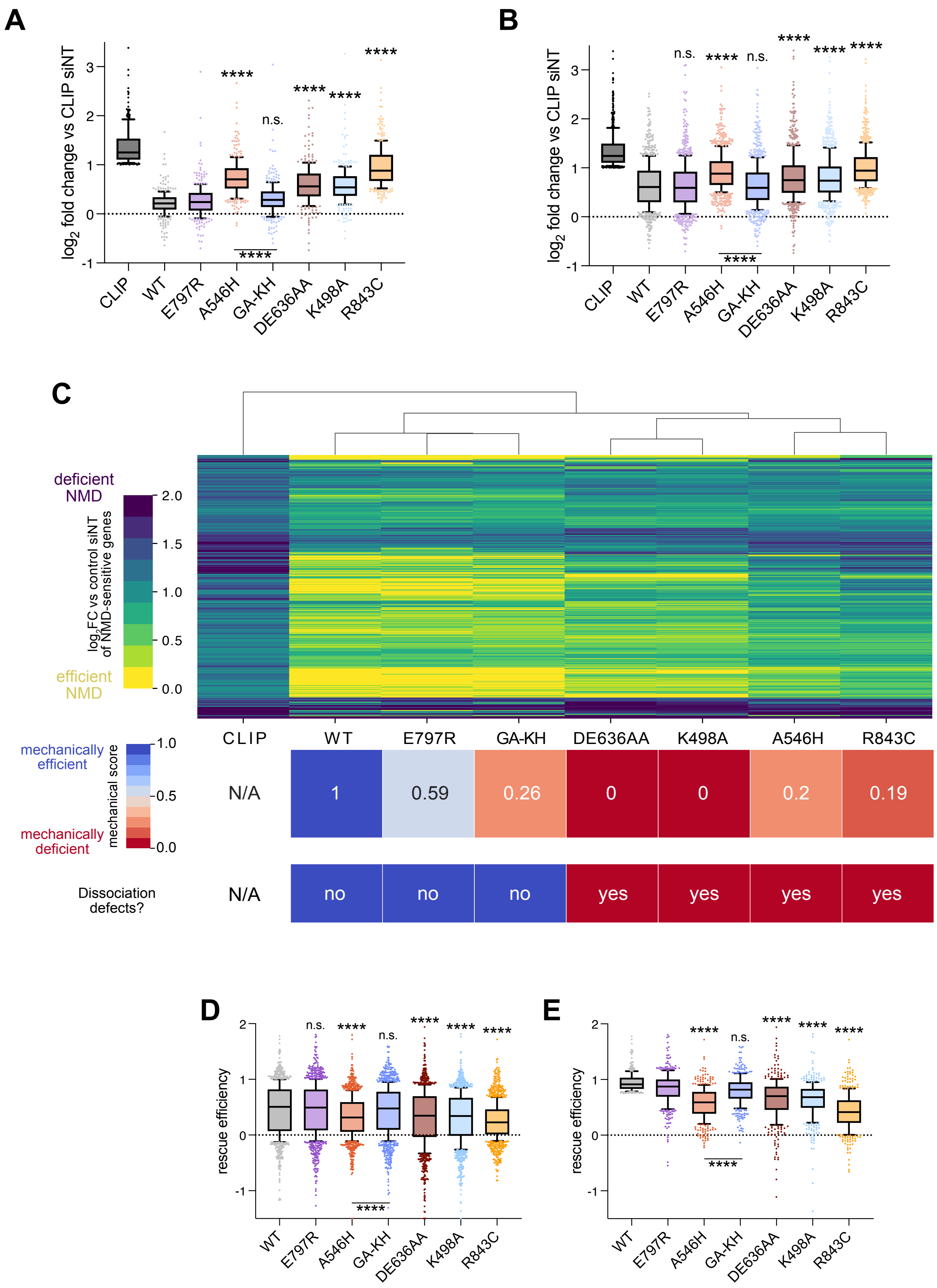


**Supplementary Figure S10: Related to Figure 5.**

***Supplementary Figure S10. RNA-seq of knockdown/rescue experiments reveals importance of UPF1 ATP-stimulated dissociation from RNA. Related to Figure 5.*** (**A**) As in Figure 5D but data visualized as a boxplot indicating interquartile ranges, with 10-90% intervals indicated by whiskers. Kruskal-Wallis tests with Dunn's correction for multiple comparisons were performed between E797R and other mutants, and between A546H and GA-KH. CLIP and WT were excluded from statistical analysis as their distributions were truncated due to filtering (*****FDR* < 0.0001). Only endogenous UPF1 knockdown conditions are shown. (**B**) As in (**A**) but only filtering for genes that were upregulated > 2-fold in the siUPF1 CLIP condition. Kruskal-Wallis statistical tests were performed between WT and the indicated conditions, and between A546H and GA-KH. CLIP was excluded from statistical analysis as the distribution was truncated due to filtering (*****FDR* < 0.0001). (**C**) As in Figure 5D but only filtering for genes that were upregulated > 2-fold in the siUPF1 CLIP condition. Second and third rows are summaries of the biochemical/biophysical characterization of UPF1-HD constructs from previous *in vitro* work (31). Mechanical scores were calculated using processivities, unwinding rates, ATPase rates, mechanochemical coupling, and protein displacement activities normalized to the WT UPF1-HD construct. Third row refers to *in vitro* ATP-dependent dissociation defects, if any. (**D**) Box plots of rMATS analysis (59), indicating interquartile ranges, with whiskers denoting 10-90% ranges. Kruskal-Wallis tests were performed between WT and the indicated conditions, and between A546H and GA-KH. CLIP was excluded from statistical analysis as the distribution was truncated due to filtering of transcripts that increased in inclusion by 10% in siUPF1 CLIP (*****FDR* < 0.0001). (**E**) As in (**D**) but only using transcripts that were rescued > 75% in the WT condition. Kruskal-Wallis tests were performed between E797R and other conditions, and between A546H and GA-KH. CLIP and WT were excluded from statistical analysis as their distributions were truncated due to filtering (*****FDR* < 0.0001). See **Supplementary Table S8** for raw data.

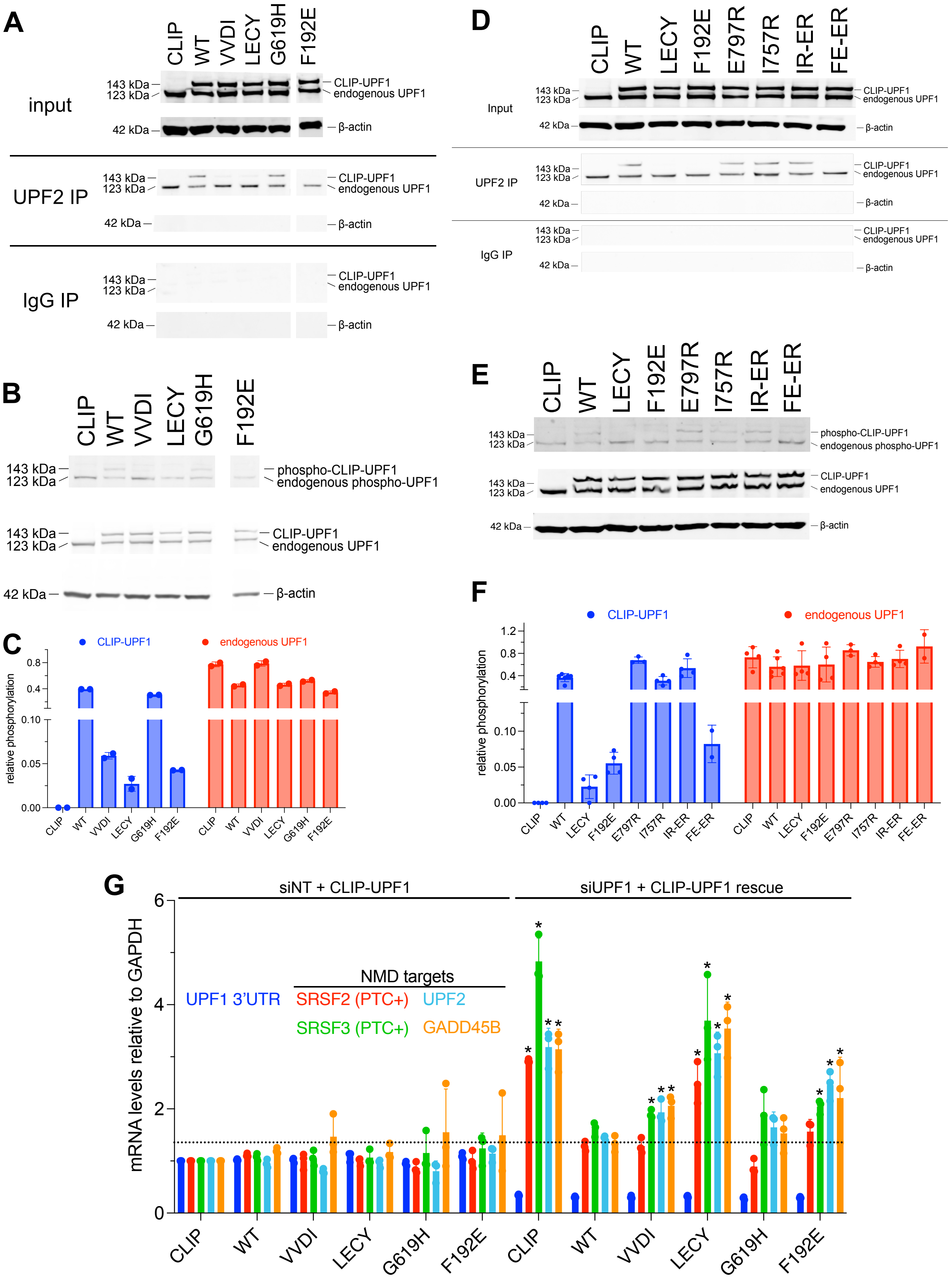


**Supplementary Figure S11: Related to Figure 5-6.**

***Supplementary Figure S11. The F192E mutation reduces UPF2 binding and steady-state phosphorylation in mammalian cells. Related to Figure 5-6.*** (**A**) Western blots of UPF1 (top blots of each condition) and β-actin as a loading control (bottom blots of each condition) after immunoprecipitation of UPF2 (UPF2 IP blots) or IgG as a control (IgG IP). Input blots indicate the input controls that were not subjected to immunoprecipitation. VVDI is VV204-5DI and LECY is LECY181-4VRVD. White space indicates different blots. (**B**) Western blots of phospho-UPF1 (top blots) using total UPF1 (middle blots) as a control and β-actin as a loading control (bottom blots). (**C**) Quantification of (**B**) from 2 independent experiments. (**D-F**) As in (**A-C**) but with a different set of CLIP-UPF1 mutants from 3 independent experiments. (**G**) mRNA levels of NMD targets relative to GAPDH using RT-qPCR normalized to the siNT CLIP condition from 3 independent experiments. UPF1 3′ UTR mRNA levels were measured to determine the extent of endogenous UPF1 knockdown. SRSF2 (PTC+) and SRSF3 (PTC+) primers amplify the PTC-containing exon. Error bars represent standard deviation. siNT is non-targeting siRNA and siUPF1 is UPF1 siRNA. A two-way ANOVA statistical test was performed, with comparison between each siUPF1 condition and the siNT CLIP condition (*$P < 0.001$, **Supplementary Table S7**).
